# The multi-level regulation of clownfish metamorphosis by thyroid hormones

**DOI:** 10.1101/2022.03.04.482938

**Authors:** Natacha Roux, Saori Miura, Mélanie Dussene, Yuki Tara, Fiona Lee, Simon de Bernard, Mathieu Reynaud, Pauline Salis, Agneesh Barua, Abdelhay Boulahtouf, Patrick Balaguer, Karine Gauthier, David Lecchini, Yann Gibert, Laurence Besseau, Vincent Laudet

## Abstract

Most marine organisms have a biphasic life cycle during which a pelagic larva is transformed into a radically different juvenile. In vertebrates the role of thyroid hormones (TH) in triggering this transition is well known, but how the morphological and physiological changes are integrated in a coherent way with the ecological transition remains poorly explored. To gain insight into this question, we performed an integrative analysis of metamorphosis of a marine teleost, the clownfish *Amphiprion ocellaris*. We reveal how TH coordinate a change in color vision as well as a major metabolic shift in energy production, hence highlighting its central integrative role in regulating this transformation. By manipulating the activity of LXR, a major regulator of metabolism, we also reveal a tight link between metabolic changes and metamorphosis progression. Strikingly, we observed that these regulations are at play in the wild revealing how hormones coordinate energy needs with available resources during life cycle.

## Introduction

In vertebrates, metamorphosis is a common post-embryonic transition regulated by thyroid hormones (TH) during which a larva is transformed into a juvenile. The role of TH as the main trigger and coordinator of the distinct biological processes that occurred during metamorphosis has been extensively studied, mostly in anurans as well as in flatfish (Power et al., 2008; Tata, 2006). Both biological models exhibit a spectacular metamorphosis during which major morphological, physiological and ecological transitions occurred (Laudet, 2011). TH not only trigger but also coordinate at the tissue and cellular level the transformation of the larva into a juvenile. For example, in tadpole, cell-specific actions of TH are instrumental for the transformation of several organs (intestine, limbs, tail) illustrating the pleiotropic action of this hormone (Grimaldi et al., 2013). This action is mediated by the binding of TH to thyroid hormone receptors (TR) that act as transcription factor and regulate the expression of target genes (Sachs and Buchholz, 2017). However, these models (flatfish and amphibian) are more exceptions than the rule as in most vertebrate species morphological changes are more subtle (Buchholz, 2015; Holzer and Laudet, 2013; Laudet, 2011). Indeed, all chordates, including amniotes, are passing through such a post-embryonic transformative period even if, in many species, it is not morphologically spectacular (Holzer and Laudet, 2013). In addition, several reports suggest that the successful completion of metamorphosis is decisive for the quality and therefore the ecological success of the juvenile emanating from it (Besson et al., 2020). Therefore, understanding how TH control this transformation is critically important to better understand the pleiotropic action of the hormone and how juvenile can cope with their new environment (Lowe et al., 2021).

A particular challenge, common to all vertebrate species, is to make sure that the transformation is well aligned with available environmental conditions and with the metabolic status of the organism (Hulbert and Else, 2000; Sheridan and Kao, 1998). THs, with their well-known metabolic effects, are critical for this action (Sayre and Lechleiter, 2012; Sinha et al., 2018). This has been shown in several species; particularly in sticklebacks in which they control metabolic rate in freshwater populations that are living in an energy-poor environment (Kitano et al., 2010). Similarly, studies in several teleost fish have shown that TH impact the activity of enzymes involved in lipid metabolism therefore influencing energy availability (Deal and Volkoff, 2020). However, these metabolic actions of TH are mostly studied in juveniles or adults and how this occurred during metamorphosis remain largely unknown (Deal and Volkoff, 2020). Shedding light on how TH control metabolism during metamorphosis would be particularly desirable as it is known that metabolic pathways are heavily modified during larval transformation (Darias et al., 2008). Therefore, it remains to be determined: (i) how TH regulate metabolism during metamorphosis and (ii) how this action is integrated with the transformation of the larvae into a juvenile, also controlled by the hormone. To fulfil this gap, we carried out for the first time an integrative study of metamorphosis to better understand the central role of TH in the coordination of larval transformation with metabolic status. Understanding these processes is critical to better understand the ecological function of TH but also the constrains and bias affecting metamorphosis, a key process ensuring natural population replenishment. This is particularly important in the context of the global changes affecting natural populations worldwide (Lowe et al., 2021).

We used the false clownfish *A. ocellaris* as a model system to explore the hormonal basis of this metabolic integration. These fishes live in coral reefs, in symbiosis with sea anemones in which they form colonies with a reproductively active pair and a variable number of juveniles (Buston, 2003). Every two to three weeks, the couple lays ca. 500 eggs on a substrate near their host. After hatching the larvae disperse in the ocean for 10–15 days. At the end of this pelagic phase, the larvae transform via metamorphosis into small juveniles, called young recruits, which must locate a reef and actively look for a host sea anemone using their sensory abilities combining visual, chemical and acoustic cues (Barth et al., 2015). During metamorphosis, the larvae lose their larval characteristics (light pigmentation, elongated body shape) and transform into miniature adults, with an ovoid body shape and a conspicuous pigmentation pattern displaying white bars on a bright orange background (Roux et al., 2019). These pigmentation changes have been shown to be regulated by TH indicating that as for other teleost fishes this metamorphosis is controlled by TH (Holzer et al., 2017; Salis et al., 2021).

In this paper, we conducted an extensive study of vertebrate metamorphosis by combining developmental transcriptome assembly, TH levels measurements, behavioral observations, lipid content analysis, *in situ* observations, and functional experiments. By focusing on visual perception and metabolic changes, we uncover how TH control gene regulatory programs, as well as behavioral and physiological outputs, hence highlighting the central integrator role of TH in regulating the whole larva to juvenile transformation. Our results demonstrated that TH link metabolic regulation, morphological transformation and behavioral changes ensuring the full ecological transformation of the pelagic larvae into a benthic reef associated juvenile.

## Results

### Three distinct post-embryonic phases

The entire larval development period of *A. ocellaris* have been divided into seven distinct developmental stages (named S1 to S7 for stage 1 to 7 in Fig. 1A, Roux et al. 2019). To obtain a global perspective on gene expression levels during post-embryonic development, we performed a transcriptomic analysis of these seven stages. RNAs from three entire larvae per stage were extracted and sequenced on an Illumina platform giving a total of 21 samples (see Material and Methods).

**Figure 1:**
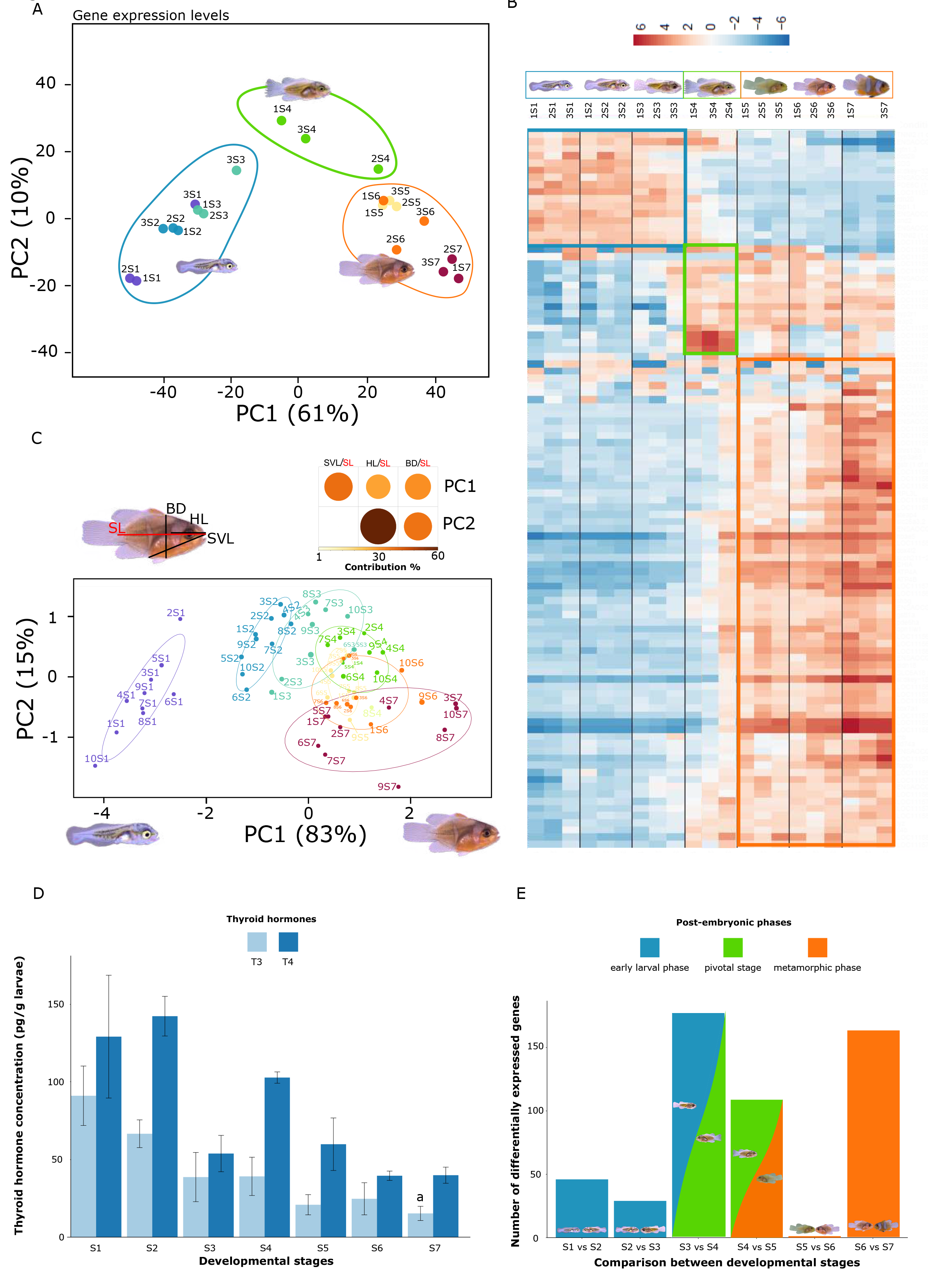
Three distinct periods during clownfish post-embryonic development. (A) Principal component analysis (PCA) according to the 1000 genes with the highest variance showing three major groups: early developmental stages, stage 4, and late developmental stages. (B) Heatmap of the 100 genes with the highest variance. Colors represent the intensity of the centered (but unscaled) signal that, for each gene, ranges from low (blue), to medium (white) and to high (red). (C) PCA of the morphological transformation according to body depth (BD), head length (HL) and Snout vent length (SVL) over Standard length (SL, in red). (D) Variation in TH levels during the seven post embryonic stages of *A. ocellaris.* (E) Number of genes differentally expressed between contiguous developmental stages.

Principal component analysis (PCA) and global hierarchical clustering were first applied on the 1000 genes with the highest amplitude of expression (variance) without considering the developmental stages (Fig. 1A and Supp. Fig. 1A). Both methods allowed distinguishing three distinct groups: (i) early developmental stages which included stages 1, 2 and 3, (ii) late developmental stages including stages 5, 6 and 7, and (iii) the pivotal stage 4. Of note, one stage 4 individual is tending toward the late stages cluster, indicating that there may be a gap between transcriptional regulation and the observed morphological changes (Fig. 1A and Supp. Fig. 1A).

The same three groups were also observed on the heat map clustering the 100 genes with the highest variance (Fig. 1B). One group showed genes that are up regulated (in red) during early larval development, corresponding to stages 1-3. Another group is formed by individuals in which genes are up regulated during late larval development (stages 5-7). Finally, some genes appeared to be up- regulated early on, at stage 4. Interestingly, when we considered the number of differentially regulated genes between each consecutive stage, we observed that stage 4 coincides with a major transition of the regulatory program with 178 significantly differentially expressed between stage 3 and 4 (Fig. 1E, Supp. Fig. 2).

**Figure 2:**
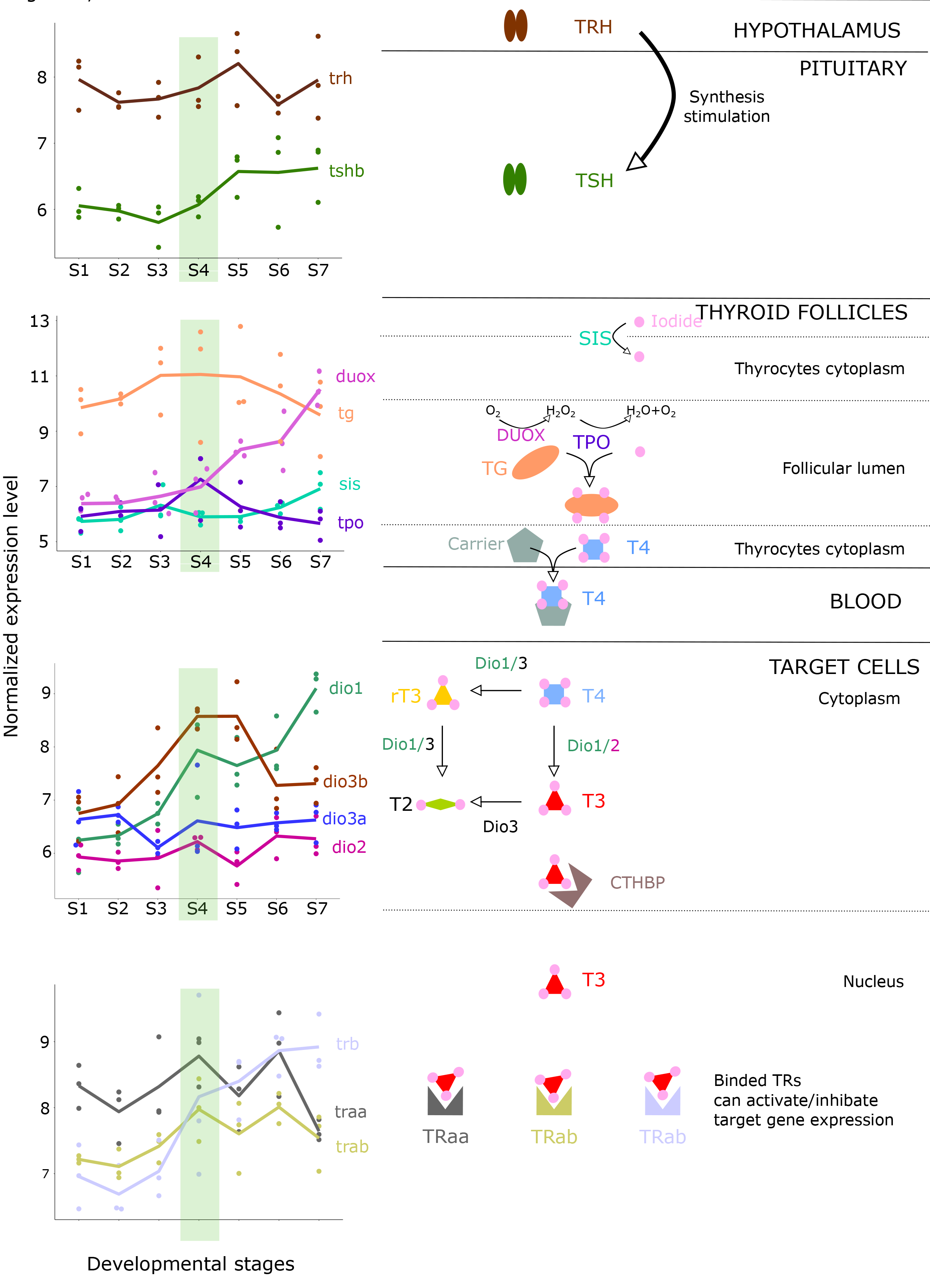
TH pathway is active during metamorphosis. Expression levels of TH signaling pathway genes extracted from the transcriptomic data, depicted in front of each step of the pathway. TRH: thyroid releasing hormone, TSH: thyroid stimulating hormone, DUOX: dual oxidase, TG: thyroglobulin, TPO: thyroperoxidase, NIS: sodium iodine symporter, DIO: deiodinase, TR: thyroid hormone receptor.

The two periods we observed are distinct in terms of morphological transformation. On a PCA performed with morphological measurements (body depth (BD)/standard length (SL); head length (HL)/SL and snout vent length (SVL)/SL), we observed that the three first stages are aligned along the main axis of variation (PC1), whereas after stage 4 there is an inflexion and the metamorphic stages are, then, varying mostly along PC2 (Fig. 1C). These changes can be explained by a change in body shape, metamorphic stages becoming more ovoid (increase of body depth). During the last stages, we also observed major changes in body pigmentation. We conclude that stages S5 to S7 correspond to the metamorphosis *per se,* whereas stage 4 appears to be a pivotal stage assuring the transition between the larval period and the metamorphosis.

Interestingly, TH levels measurements showed a high level of T4 and T3 at stage 1 and 2 that likely correspond to residues of maternal loading as observed in other species (Fig. 1D) (see for example (Chang et al., 2012; Einarsdóttir et al., 2006a) on Atlantic halibut and Chang et al., 2012; on zebrafish). Then, we observed a surge of T4 levels at stage 4 which precedes white bars appearance occurring at stage 5 (Roux et al. 2019; Salis et al., 2020) followed by a decline in later stages. T3, the most active form of TH, showed on the contrary more stable levels than T4, probably linked to the fact that T3 is mostly produced and metabolized by deiodinases with different kinetics in different peripheral tissues (Darras and Van Herck, 2012).

Overall, these results reveal that *A. ocellaris* post-embryonic development is characterized by three distinct phases: (i) larval development, (ii) the pivotal stage 4 that marks the onset of metamorphosis with a peak of TH and (iii) metamorphosis that corresponds to the actual transformation.

### TH pathway is activated during metamorphosis

To investigate the role played by TH in this process, we analyzed the expression levels of the TH signaling genes. We selected genes from the hypothalamo pituitary thyroid (HPT) axis involved in the central control of TH synthesis (*trh:* thyrotropin releasing hormone, and *tshβ:* thyrotropin stimulating hormone subunit *β*) ; genes involved in TH synthesis in the thyroid follicles (*tg: thyroglobulin*, *tpo:* thyroperoxidase, *nis* sodium/iodine symporter, *duox:* dual oxidase); genes involved in TH metabolism (*dio1*, *dio2*, *dio3a* and *dio3b:* deiodinase 1, 2, 3a, 3b respectively); as well as the genes encoding thyroid hormone receptors (*TRαa*, *TRαb* and *TRβ* ; Fig.2).

The expression level of *trh* is relatively stable and *tshβ* expression levels rise from the onset of metamorphosis (stage 4) and stabilize until stage 7. In the thyroid gland, *duox* have been shown to be critical for TH production and is associated with high TH levels in clownfish (Salis et al., 2021). Interestingly, we observed that its expression levels strongly rise during metamorphosis, starting at stage 5. The expression level of *nis* (encoding for iodine transporter in thyroid follicles) increased, at stage 6. In contrast, the expression of the gene encoding for the precursor of TH, *tg,* increased early on (stage 3) and reached a peak at stage 4 as did the expression of *tpo*. These results clearly corroborate the high production of T4 observed in stage 4 confirming that this corresponds to the onset of metamorphosis.

Deiodinases are known to be key enzymes controlling the amount of T3, the most active thyroid hormone, available in peripheral tissues and their expression in specific tissues are known to be excellent indicators of the activity of the pathway (Bianco and Kim, 2006). We observed a spectacular peak of expression of *dio3b*, one of the duplicates of the *dio3* gene which is coherent with the known regulation of this gene by high TH levels (Russo et al., 2021). We also observed a strong two step increase of *dio1* (known to either activate or inactivate TH depending on the context) at stage 4 and stage 7, respectively. In contrast, both *dio3a* and *dio2* (which activates TH by converting T4 into T3) expression remains constant. This lack of change in global expression levels for *dio3a* and *dio2* may hide significant differences at tissues levels that are not visible in the context of entire larvae as it is also the case for the global T3 levels.

Finally, we analyzed the expression levels of the three TR that are known to mediate the action of TH in target cells: *TRβ* displayed a spectacular increase of expression starting at stage 4, clearly in accordance with its role as a TH regulated gene in many species (McMenamin and Parichy, 2013). Its expression level remains high until stage 7. *TRαa* and *TRαb* expression peaked at the stage 4 and stage 6. The statistical significance of the variations of expression levels described above are summarized in Supp. Fig. 1B. A global PCA analysis of TH signaling gene expression level again separated the three first stages from S5-S7, indicating a clear activation of this pathway at stage 4 (Supp. Fig. 1C). Taken together, TH signaling gene expression levels are in accordance with the peak of T4 observed at stage 4 confirming that clownfish metamorphosis starts at this stage.

The above results suggest that several of these TH signaling genes may be regulated by TH as observed in other species (Tata 2006; Power et al. 2008). Additionaly, after investigating the effects of exogenous TH by treating stage 3 larvae with three different concentrations of T3 we observed a positive or negative regulation of TH signaling genes. This is very similar to what has been observed in other species (Campinho, 2019; McMenamin and Parichy, 2013). In summary, our analysis clearly suggests that TH-signaling is activated during clownfish metamorphosis and that TH are the main driver of the complex transformation of a pelagic larva to a reef associated juvenile.

### TH control a molecular and behavioral shift in vision

During their transformation, clownfish larvae must switch from an oceanic (pelagic) environment to a colorful reef environment. It is well known that in many fish species this ecological transition is accompanied by a change in color vision (Cortesi et al., 2016). Since TH appeared critical for larval to juvenile transition in clownfish, we investigated the regulation of genes encoding for visual opsin during metamorphosis.

Eight visual opsin genes (*opnsw1-α, opnsw1-β, opnsw2B, rh1, Rh2A-1, rh2A-2, rh2B, opnlw)* have been identified in *A. ocellaris* (Mitchell et al., 2021). In our transcriptomic data we observed a reciprocal shift in their expression (Fig. 3A): short wavelength opsins (*opnsw1-α, opnsw1-β, opnsw2B*), and mid-wavelength opsins (r*h2A, rh2B*) are highly expressed at the beginning of larval development and are down-regulated after stage 4. Of note we observed a larval expression of both duplicates of *opnsw1*, including *opnsw1-α* for which no clear expression was yet detected (Mitchell et al., 2021). In contrast, the long wavelength opsin (*opnlw*) is poorly expressed in larval stages and its expression strongly increase from stage 4 onwards and remains high throughout metamorphosis (Fig. 3A).

**Figure 3:**
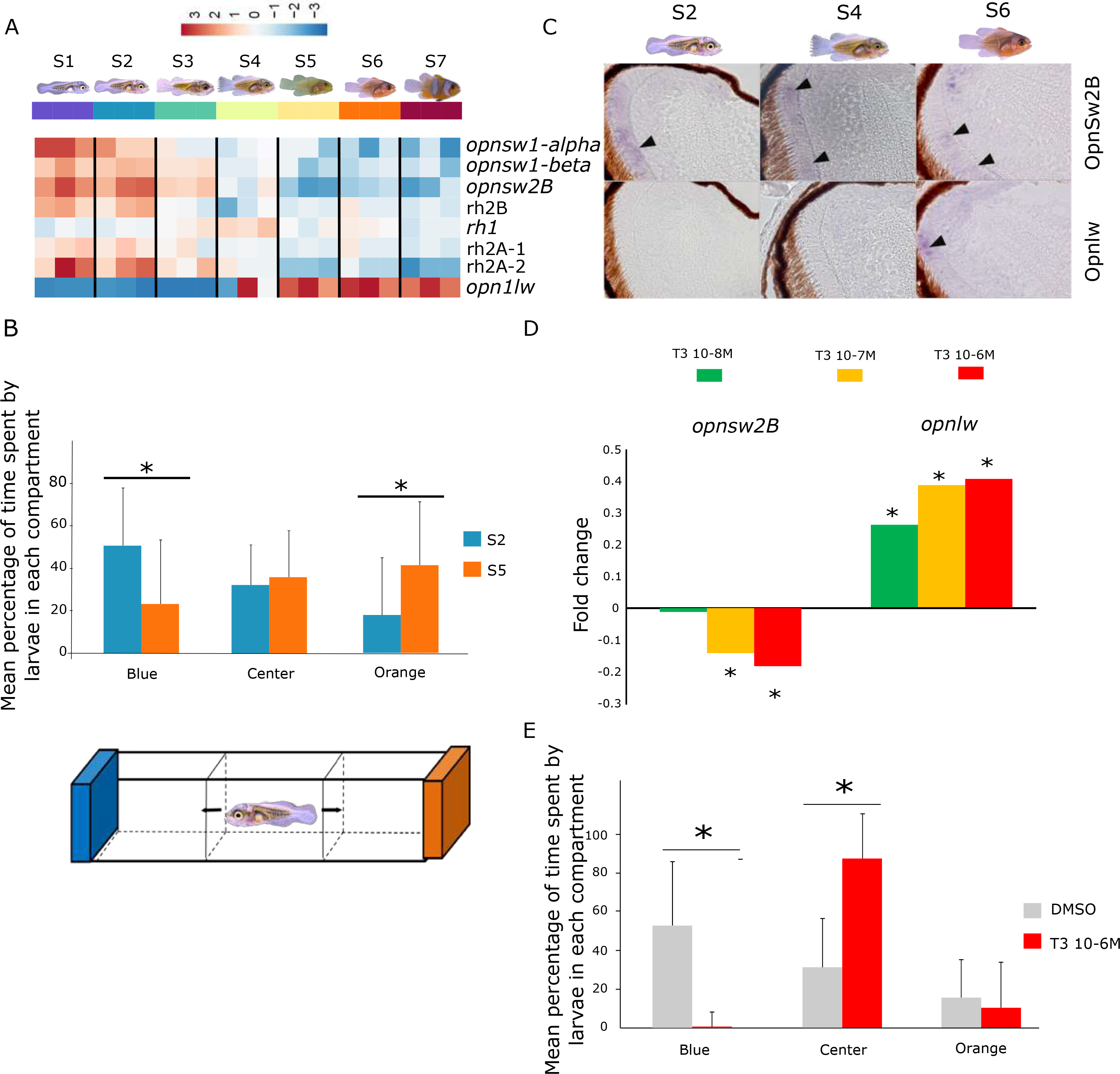
TH control visual perception shift by regulating opsin gene expression. (A) Heat map of the eight opsin genes involved in visual perception organized from top to bottom according to short (*opnsw2B, opnsw1α, opnsw1β*), medium (*rh2B, rh2A-1, rh2A-2, rh1*) and long (*opnlw*) wavelength sensitivity. (B) Behavioral test for color preference. Mean percentage of time spent by stages 2 (n=20) and 5 (n=20) larvae in each compartment of a dual choice chamber. Statistical differences (indicated by a star (*)) between each stage in each compartment were calculated by a chi² test. (C) *In situ* hybridization of sectioned stage 2, 4 and 6 larval retina using probes for *opnsw2b* and *opnlw*. Black arrowhead indicates expression signals in the photoreceptors (external nuclear layer). (D) Regulation of *opnsw2B* and *opnlw* expression after 48 hours of treatment at 10^-8^, 10^-7^, 10^-6^ M T3 + IOP (n=3 pools of 2 larvae per condition). Expression levels are expressed in fold change and statistical differences between treatment and DMSO control are indicated by a star (*). (E) T3 affect the color preference. Effects of T3 at 10^-6^M (n=19) compared to DMSO control (n=14) after 48 hours treatment on the mean percentage of time spent by larvae in each compartment. Statistical differences (indicated by a star (*)) between each condition in each compartment were calculated by a chi² test.

This reciprocal shift (decrease in shortwave length and increase in long wavelength opsins, respectively) suggested that there could be a shift in visual perception from blue/green to yellow/red during metamorphosis. We therefore tested the visual preferences of *A. ocellaris* before and after metamorphosis using a dual choice chamber (illustrated in Fig. 3B). Stage 2 larva spent significantly more time in the blue compartment (50% vs. 23% in the orange compartment *p-value* = 0.004) whereas stage 5 larva prefer the orange one (41% vs. 17% in the blue compartment, *p-value* = 0.009). This indicated that opsin gene expression correlates with visual preference. This shift is also visible by *in situ* hybridization: The *opnsw2B* gene is strongly expressed in photoreceptors at stage 2 and slightly visible at stage 4 and 6 on the perimeter of the retina except on the ventral side whereas the *opnlw* gene is not expressed at stage 2 and is detected in the photoreceptors at stage 6 (Fig. 3C).

We then tested if TH is controlling this molecular and behavioral shift. We observed that T3 strongly up regulated the expression levels of *opnlw* and down regulated *opnsw2B* from 12 to 72 hours after treatment (Fig. 3D, Supp. Fig. 4A). It also up regulated the expression of *Rh2A* after 48 hours of exposure (Supp. Fig. 4A). These data therefore suggest that the shift in opsin gene expression is controlled by TH.

**Figure 4:**
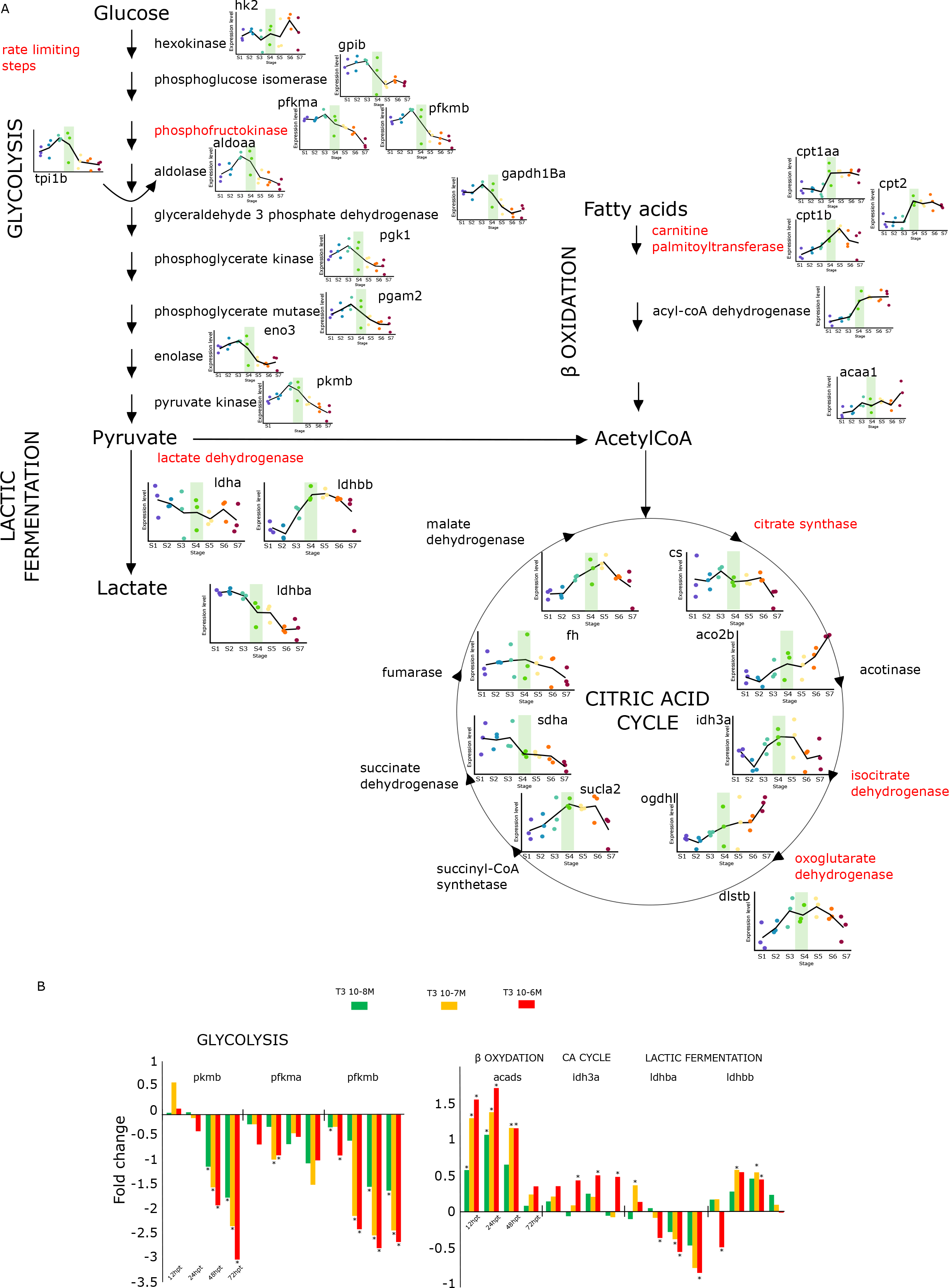
TH induce a major metabolic shift during metamorphosis. (A) Expression levels of genes involved in glycolysis, lactic fermentation, fatty acid β oxidation, and citric acid cycle at each developmental stage (S1: stage 1 ; S2: stage 2 ; S3: stage 3 ; S4: stage 4 ; S5: Stage 5 ; S6: stage 6 and S7: stage 7) extracted from transcriptomic data clearly revealing a change of expression at stage 4 (indicated on each graph by a green rectangle) for each pathway. (B) Effects of T3 (10^-8^, 10^-7^, 10^-6^ M, n=3 pools of 2 larvae per condition) after 12-, 24-, 48- and 72-hours post treatment (hpt) on the expression levels of the representative genes for glycolysis (*pkmb, pfkma and pfkmb),* β oxidation (*acads*), citric acid cycle (*idh3a*), lactic fermentation (*ldhba, ldhbb*).

We next investigated if TH could also induce a shift in visual preferences. We treated stage 3 larvae (*i.e.,* before metamorphosis) for 72 hours with either T3 or DMSO as a vehicle and measured the color preference of the larvae in a dual chamber, as described above. Interestingly, we observed that control larvae spent significantly more time in the blue compartment than T3 treated larvae (53% vs 1% respectively, *p-value*=5,2x10^-7^, Fig. 3E), which can be correlated with the higher expression of *opnsw2B* observed in control larvae (Fig. 3D). Surprisingly, T3 treated larvae did not spend much time in the orange compartment compared to DMSO treated larvae (11% vs 16% respectively). Instead, they remained most of the time in the central compartment (88% vs 31% for the DMSO controls, *p-value*=1,3x10^-5^). Observation of the larvae after 72 hours of T3 treatment during the experimental trial revealed however that they remained close to the bottom of the chamber, barely swimming, whereas control larvae were actively exploring the blue compartment. The same behavior was observed when 30 days post hatching (dph) juveniles were similarly tested (Supp. Fig. 4B). This indicated that, as expected, T3 accelerates the appearance of a juvenile-type behavior which is in accordance with the benthic lifestyle of juvenile’s clownfish, hiding in the tentacles of their sea anemone host.

Taken together these results clearly demonstrate that TH control a shift in opsin gene expression that coincides with a change in color preference occurring during metamorphosis. Even if the shift of opsin gene expression cannot solely explain the visual preference, our results clearly indicate that TH is coordinating a molecular, behavioral, and ecological transition essential for larval survival in the wild.

### A TH regulated metabolic transition during metamorphosis

Because TH is known to regulate metabolism in mammals, we investigated metabolic gene expression during clownfish metamorphosis (Mullur et al., 2014). Figure 4 shows the expression profile of the main genes involved in these pathways and highlights the rate limiting steps for glycolysis (phosphofructokinase, *pfkma* and *pfkmb*), citric acid cycle (citrate synthase, *cs*; isocitrate dehydrogenase, *idh3a*; oxoglutarate dehydrogenase, *ogdhl*, *dlst2*) and fatty acid β-oxidation (carnitine palmitoyl transferase *cpt1aa*, *cpt1b*, *cpt2*). The expression profile of all the genes implicated in these pathways are shown in Supplementary figures 5, 6 and 7.

**Figure 5:**
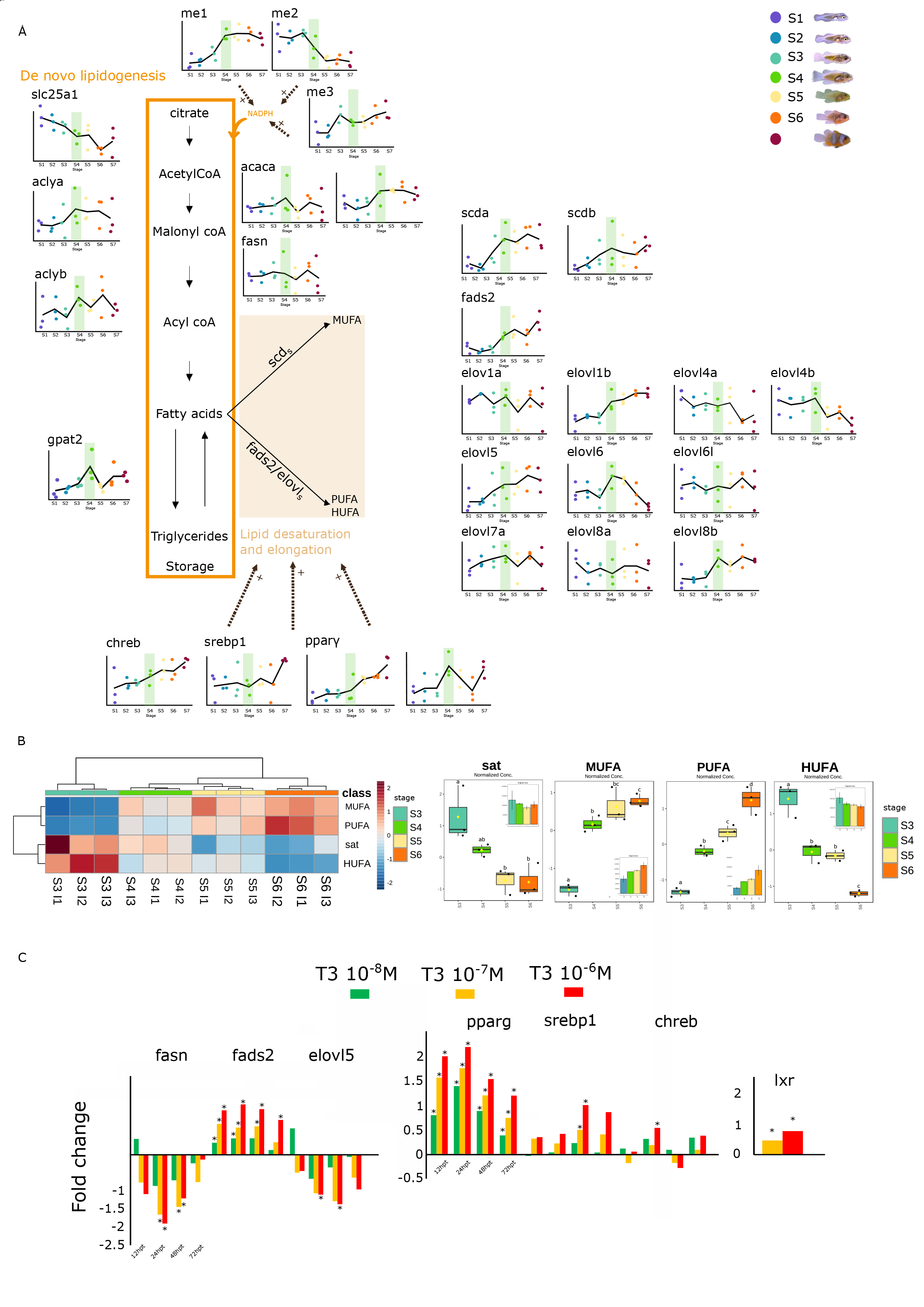
TH action on lipid metabolism regulation during *A. ocellaris* metamorphosis. (A) expression levels of genes involved in *de novo* lipogenesis and lipid desaturation and elongation at various stages (S1: stage 1; S2: stage 2; S3: stage 3; S4: stage 4; S5: Stage 5; S6: stage 6 and S7: stage 7). (B) Fatty acid content of *A. ocellaris* at stage 3, 4 5 and 6 illustrated with an heat map (left) and box plots (right). Significant differences between each fatty acid class are indicated by different letters. (Sat: saturated, MUFA: monounsaturated fatty acids, PUFA: polyunsaturated fatty acids, HUFA: highly unsaturated fatty acids). (C) Effects of T3 (10^-8^, 10^-7^, 10^-6^ M, n=3 pool of 2 larvae per condition) after 12, 24, 48 and 72 hours post treatment (hpt) on the expression levels of representative genes for fatty acid synthesis (*fasn)* desaturation (*fads2*), elongation (*elovl5*) and for the transcription factors involved in lipid metabolism regulation (*pparg, srebp1, chreb, lxr)*.

**Figure 6:**
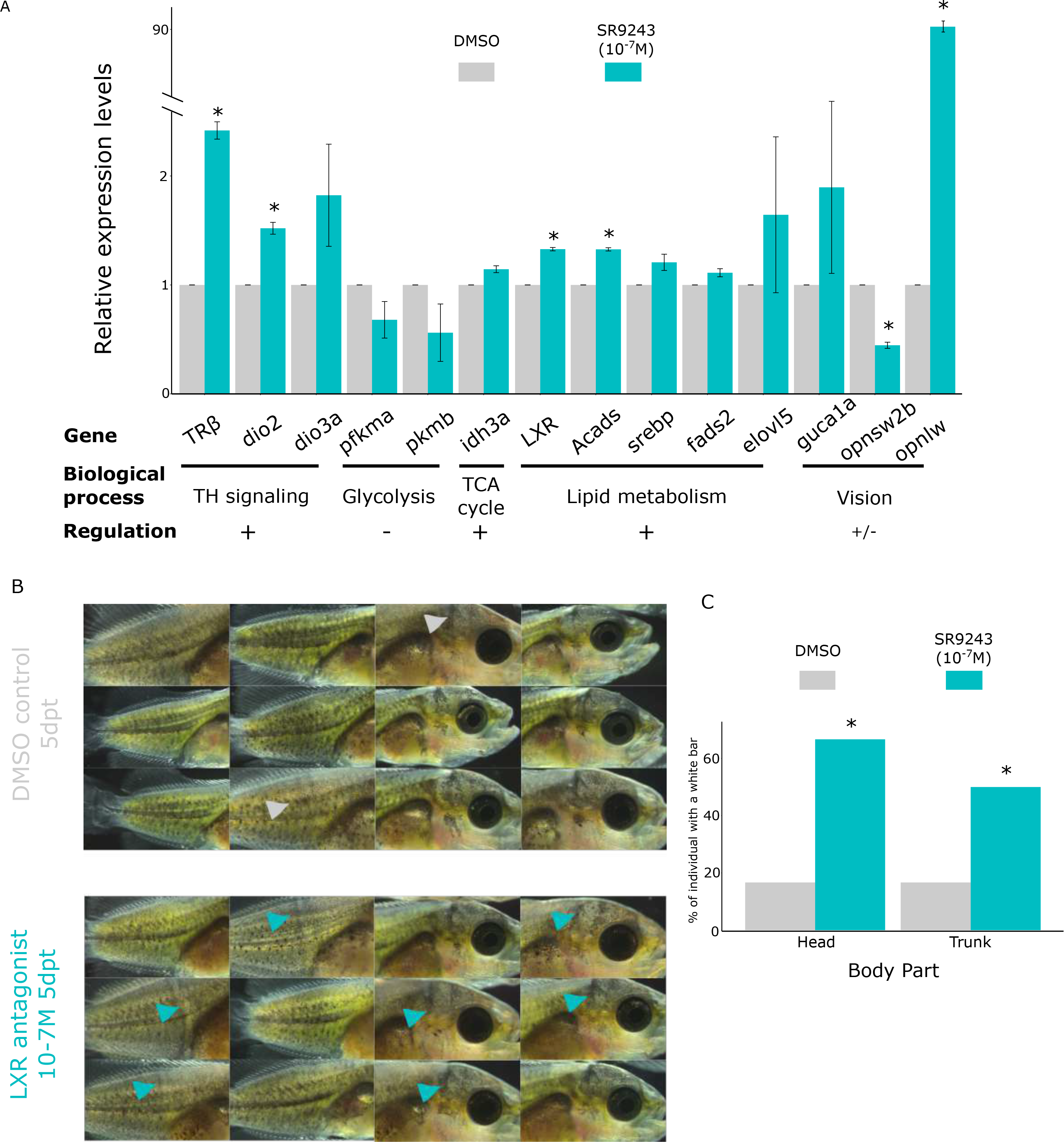
Link between metabolic changes and metamorphosis progression revealed by LXR antagonist treatment. (A) Effect of LXR antagonist at 10-7M after 5 days of treatment on the expression levels of genes involved in TH signaling (*trb, dio2, dio3a),* glycolysis (*pfkma, pkmb*), citric acid cycle (TCA) (*idh3a*), lipid metabolism (*lxr, acads, srebp1, fads2, elovl5*) and vision (*guca1a, opnsw2B and opnlw).* Significant differences are indicated by a star (*). (B) Pictures of *A. ocellaris* larvae after 5 days of treatment (dpt) with LXR antagonist (SR9243) at 10^-7^ M (bottom) compared to DMSO control (top). Grey and blue arrowhead indicates white bar appearance in controls and LXR antagonist treated fish, respectively. (C) Quantitative analysis showing the percentage of individuals presenting white bar on the head and trunk (n=6 per condition). Significant differences are indicated by a star (*).

**Figure 7:**
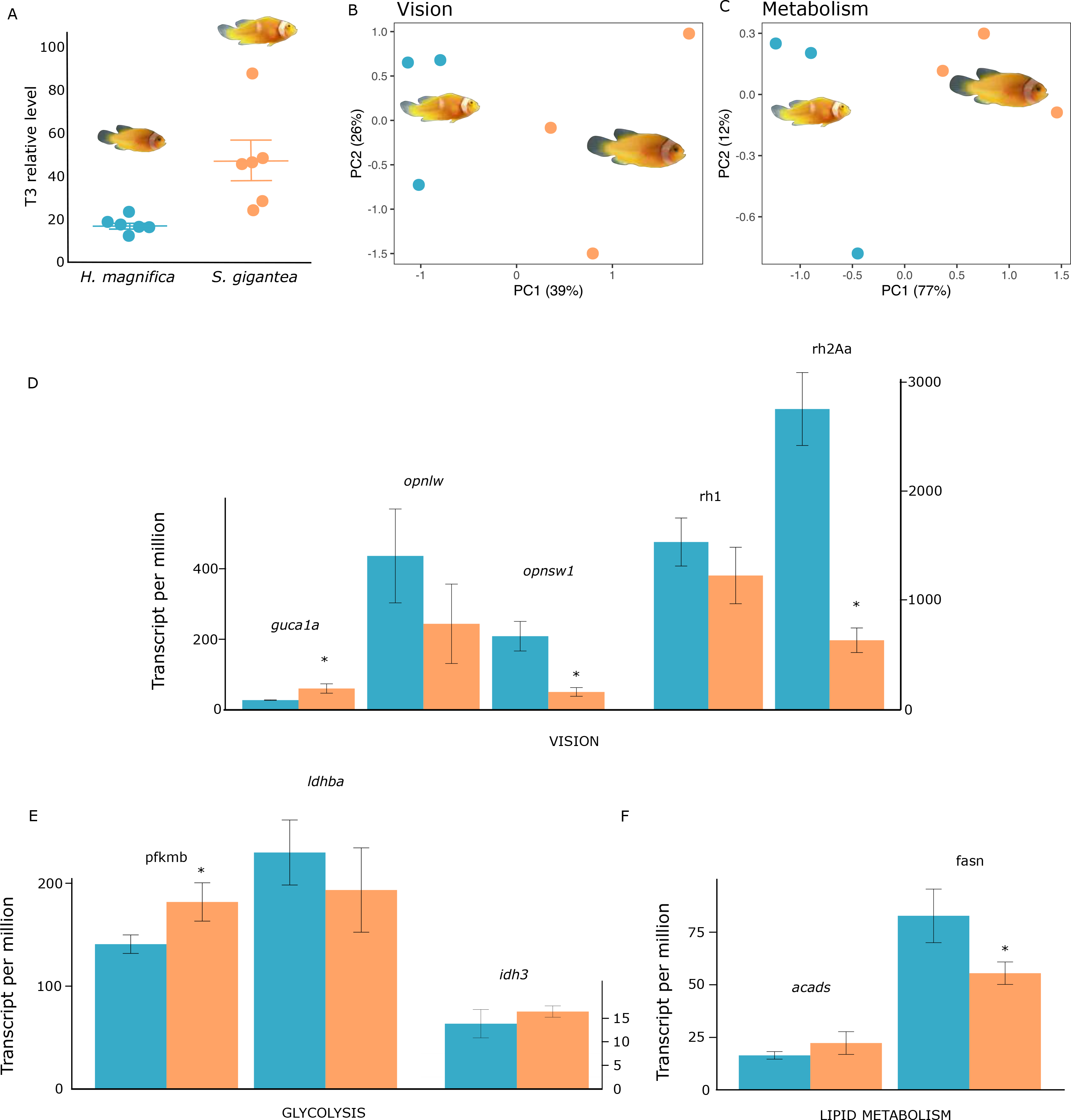
Natural variations in TH levels reveal metabolic and visual effect. (A) Thyroid hormone level (T3) of new recruits sampled in the sea anemone *Heteractis magnifica* (blue, n=5) versus *Stichodactyla gigantea* (orange, n=5) (from Salis et al., 2021). (B-C) Principal component analysis of genes involved in vision and metabolism showing the separation between *A. percula* recruits sampled in *H. magnifica* vs *S. gigantea.* (D-E-F) *E*xpression levels of genes involved in vision (D) and metabolism (E and F), obtained after RNA sequencing of *A. percula* recruit sampled in both sea anemone species, n=3 per condition.

These profiles revealed a clear overall pattern: glycolysis genes are highly expressed in larval stages (S1 to S3) while their expression decreases during metamorphosis. This is particularly true for the expression of the rate limiting enzymes (*pfkma* and *pfkmb*) which surges at S3 and strongly decreases up to stage 7. In sharp contrast, the expression of fatty acid β-oxidation genes showed an inverse profile with a low expression in larval stage and a sharp increase starting at stage 3 or 4. This is especially the case for the *acads* gene that encodes acyl-coA dehydrogenase, an enzyme involved in the second step of β-oxydation. This trend is again encountered on the rate limiting step (*cpt1aa*, *cpt1b*, *cpt2*). The citric acid cycle genes are, in overall, expressed at lower level during larval stages and their expression tends to increase early on during metamorphosis. This trend is more subtle and early than the ones observed with glycolysis and fatty acid β-oxidation but again is observed on the three rate limiting steps (*cs*, *idh3a, ogdhl* and *dlst2*). The lactic acid fermentation genes (lactate dehydrogenase, *ldha*, *ldhba*, *ldhbb*, *ldhd*) are showing a more complex regulation.

Taken together, these results suggest that larval fish mainly rely on glycolysis and lactic acid fermentation (considered as anaerobic energy production) whereas during metamorphosis, the young juveniles rely more on fatty acid, degraded by the β-oxidation, and use the citric acid cycle for aerobic energy production. Of note, this trend is the inverse to what is observed in some other fish species such as the sea bass (Darias et al., 2008; Mazurais et al., 2011). This difference is likely to be explained by a difference in life history trait, sea bass being a pelagic fish whereas clownfish is reef associated.

We then studied if TH were instrumental in controlling this transition in energy production by investigating their effects on the expression levels of *pfkma* and *pfkmb* (rate limiting step) and *pkmb* for glycolysis, *acads* for fatty acid β-oxidation, *idh3a* for the citric acid cycle as well as *ldhba* and *ldhbb* for lactic acid fermentation (Fig. 4B). In accordance with expression profile described above, TH down regulated the expression of glycolysis gene and up regulated those involved in the citric acid cycle and β-oxidation. For lactic acid fermentation we observed a dual effect with TH up regulating *ldhbb* and down regulating *ldhba*, consistent with the respective increase and decrease of those two genes during metamorphosis (Fig. 4A-B). This clearly indicates that TH control a switch between glucose-based anaerobic toward a fatty acid, aerobic-energy production.

### TH favors a complexification of lipids during metamorphosis

The previous data suggest a transition in energy source during metamorphosis with fatty acid β-oxidation being the main energy pathway during metamorphosis. In line with this observation, we noticed that the genes implicated in lipid biosynthesis and fat storage are not induced and sometimes even repressed during metamorphosis.

Lipid biosynthesis requires the transport of citrate from the mitochondria to the cytosol by the tricarboxylate carrier protein SLC25A1 and then the formation of acetyl-coA by the ATP citrate lyase (*acly*) (Fig. 5A). This acetyl-coA is then used by acetyl-coA carboxylase (*acaca* and *b*) that form malonyl-coA, the main starting susbstrate for fatty acid biosynthesis by fatty acid synthase (*fasn*). In our transcriptomic data, we observed that *slc25a1* expression decreased regularly from stage 1 until stage 6 with a final slight increase in stage 7 (Fig. 5A). The following genes in the pathway were either slightly increasing (*aclya*, *acacb*) or constant (*aclyb*, *acaca* and *fasn*). All this suggest that *de novo* fatty acid biosynthesis is minimal during metamorphosis. In accordance with this notion, TH treatment decreased *fasn*expression, suggesting that TH does not favor *de novo* fatty acid biosynthesis in this context (Fig. 5B). These data also suggest that the main source of fuel for fatty acid β-oxidation are fatty acid coming from the diet, a situation that has already been observed in many marine fish species as fatty acids are abundant in marine algae and zooplankton (Tocher, 2010).

In sharp contrast to these *de novo* fatty acid biosynthesis steps, we observed that the genes implicated in the desaturation and elongation of fatty acids are regulated during metamorphosis. This is important as long-chain fatty acid (polyunsaturated and highly unsaturated fatty acid, respectively PUFA and HUFA) participate in many biological processes and are precursors of key signaling molecules. PUFAs and HUFAS are generated by the action of front-end desaturases (FADS, SCD) and elongase (ELOVL). In contrast to mammals, in clownfish there is only one FADS that is encoded by the *fads2* gene, as in most marine fishes. This gene is up regulated during metamorphosis (Fig. 5A). The other desaturases, stearoyl-coA desaturase (*scda* and *scdb*) are also activated with similar dynamics. The many elongase genes, including *elovl5* acting on C18 and C20 PUFAs, *elovl4* also acting on PUFAs and *elovl6* acting on saturated fatty acids (FA) and monounsaturated fatty acids (MUFA) (Xie et al., 2021) showed a wide variety of patterns suggesting an intricate level of regulation: *elovl5* expression is up-regulated so are the two *elovl6* genes, whereas *elovl4* expression decreases. We also observed that *srebp1*, *pparγ*, *lxr* and *chreb*, transcription factors known to be implicated in the regulation of *fads* and *elovl* genes in fish, are also up-regulated during metamorphosis suggesting that a major coordinated change of gene expression related to lipid biochemistry occurred at this stage (Fig. 5A) (Alves Martins et al., 2012; Pinto et al., 2016; Zhang et al., 2016).

These data reveal that, while fatty acids are used as energy source during metamorphosis, they are also metabolized to generate more complex lipids that are precursors for signaling molecules. We therefore compared the lipids content between four post-embryonic stages: S3, S4 the pivotal stage, and two metamorphic stages, S5 and S6 (n=3 per stage; Fig. 5B). In accordance with the increased expression of *fads2*, *elovl5* and *elovl6*, we observed a strong statistically significant decrease of saturated lipids and, concomitantly, an increase in MUFA and PUFA. In contrast to those two fatty acid classes, HUFA content decreased steadily during metamorphosis. Supplementary Figure 8 show the complex changes of individual lipid molecules that we observed. Taken together, these results suggest that the metamorphosis coincides also with a major change in fatty acid biochemistry and the complexification of the molecules present that can be used as cell membranes constituents, energy storage, fatty acid transport and as a source of signaling molecules (Hulbert, 2021).

To determine if these changes are under TH coordination, we analyzed the effects of TH on the expression of key enzymes implicated in fatty acid desaturation and elongation and we studied if TH treatment (10^-7^ and 10^-8^ M, 72 hours post treatment of stage 3 larvae) was effectively able to alter the amount of specific lipids. At the gene expression level, we observed that TH exposure effectively induces a change in the expression of key genes, increasing *fads2* expression and, surprisingly, repressing *elovl5* gene expression (Fig. 5C). Interestingly, we noticed that TH also up regulate the expression of *pparγ*, *srebp1*, *chreb* and *lxr*, major transcriptional regulators of desaturases and elongases.

Concerning lipids, we observed a great variability from one fish to another, a situation probably linked to the fact that it is almost impossible to control the metabolic status and feeding time of such tiny aquatic organisms. This variability impacted the number of lipids for which we can get statistically significant differences in level between TH treatment and controls (DMSO). However, we observed that 37 lipids upon the 1323 studied show differences in levels after TH treatment (Supp. Fig. 9) and we noticed a non-statistically significant trend for 197 others (data not shown). In overall our data suggest that 144 lipids are increased after TH treatment whereas 137 are decreased revealing a major implication of TH in lipid metabolism.

Taken together these data suggest that during metamorphosis, clownfish used free fatty acids (saturated and HUFA) as a major energy source and, in addition, dietary lipids as substrates to generate a pleiad of molecules, many of which served as fatty acid transport (acyl carnitine), energy storage (triacylglycerol), membrane constituents (ceramide, sphingomyelin, phosphatidyl ethanolamine) or signaling molecules (phosphatidylglycerol sphingomyelin). As vision and energy metabolism, this transition in lipid biochemistry is also controlled by TH that appear as a major conductor of the metamorphosis process.

### LXR modulation reveal an intimate link between metabolic transition and metamorphosis

In the transcription factors regulating fatty acid biochemistry, LXR (liver X regulators, also called NR1H3) is particularly interesting given its known pivotal role. This receptor is a major regulator of lipid metabolism and also regulates fatty acid metabolism through, for example, the control of *srebp* and elongases and desaturases genes (Wang and Tontonoz, 2018). Additionally, in mouse *lxrβ* control TH signaling in the brain and adipose tissue and its knock-out resulted in increased *dio2*, *nis* and TH transport genes (Ghaddab-Zroud et al., 2014; Miao et al., 2015). We used SR9243, a selective LXR antagonist to test if inhibiting LXR action will affect clownfish metamorphosis and its underlying gene regulatory program. Importantly, we verified that this compound specifically inhibits fish LXR activity and is inactive on PPAR*γ* and PXR (Supp. Fig. 10).

As expected, after treating stage 3 larvae for 5 days with 10^-7^M SR9243, we observed a dysregulation of lipid metabolism genes such as *acads, srebp, fads2* and *elovl5* genes (Fig. 6A) confirming that in clownfish as in zebrafish, LXR is a regulator of lipid metabolism. We also observed that glycolysis genes (*pfkma* and *pkmb*) are down regulated whereas citric acid cycle genes are up regulated (Fig. 6A).

Interestingly, in this context, we also observed a clear acceleration of metamorphosis revealed by white bar appearance (Salis et al., 2021; Figure 6B-C). By counting the number of individuals harboring white bars we observed a significant difference in SR9243-treated fishes (66% for head bar and 50% for trunk bar vs 16% and 16% in DMSO controls, respectively; Fig. 6C). This shows that, as in mouse, LXR inhibition activate TH signaling and we effectively observed that *trβ*, *dio2* and *dio3a* are up regulated after treatment (Fig. 6A).

Based on these results, we investigated if SR9243 exposure also affects the other processes that are coordinated by TH and in particular vision. Interestingly, LXR antagonist also promoted the opsin genes expression shift, decreasing short wavelength and increasing long wavelength opsin expression (Fig. 6A). LXR antagonist also regulated the phototransduction gene *guca1a,* as observed in zebrafish (Fig. 6A ; Pinto et al., 2016). Taken together, these results show that modulating lipid homeostasis by inhibiting LXR activity affect TH signaling and vision, linking metabolic regulation with the coordination of metamorphosis.

### Natural TH regulation in an ecological context, reveals metabolic and visual effect

We previously reported a natural situation in which endogenous TH level regulation occurs (Salis et al., 2021). In Kimbe bay, Papua New Guinea *Amphiprion percula* can inhabit in two different sea anemone hosts: the carpet sea anemone *Stichodactyla gigantea* and the magnificent sea anemone *Heteractis magnifica*. In these two hosts, the new recruits exhibit differences in white bar formation that are linked to a higher TH level (Fig. 7A) and higher *duox* expression in fish hosted by *S. gigantea* (Salis et al., 2021). Given the results described above, we therefore measured the expression levels of genes implicated in vision and metabolic regulation in this natural context.

We compared gene expression between new recruits found in *H. magnifica* (n = 3) or *S. gigantea* (n = 3) by RNA sequencing of whole fish. On a PCA analysis based on vision and metabolic genes, we observed a clear separation between both types of recruits (Fig. 7B). We noticed that, in accordance with the higher TH level in *S. gigantea* recruits, the phototransduction gene *guca1a* is up regulated whereas the short wavelength opsin *opnsw1* as well as the rhodopsin *rh1* and *rh2a* are down regulated (Fig. 7D). We also detected a small down regulation of the long wavelength opsin *opnlw*.

At the metabolic level we also observed a clear separation of recruits living in *S. gigantea vs.* those in *H. magnifica* suggesting that effectively the metabolic status of both type is different (Fig. 7C). When individual genes are studied, we observed that *S. gigantea* recruits contain higher TH level, a decrease in expression of the glycolytic *pkmb* gene and the lactic fermentation gene *ldhba* and an increased expression of the TCA cycle gene *idh3* and the β-oxidation genes *acaa1* and *acads* (Fig. 7E-F). Similarly, we observed a decrease in the expression level of the fatty acid synthase gene *fasn*. These results are in accordance with the metabolic transition observed during metamorphosis and demonstrate the relevance of these effects in natural populations living in their pristine environment.

## Discussion

In this paper we reveal how TH control and coordinate a major ecological transition, the metamorphosis of a pelagic coral reef fish larvae into a benthic reef associated juvenile. Our data therefore suggest a model, discussed below, according to which TH has a central coordinating function to ensure the ecological success of the metamorphosed juvenile.

Three main functions were attributed to TH in vertebrates: (i) the triggering of metamorphosis in amphibians and fish; (ii) the regulation of adult physiology and metabolism in mammals and (iii) the adaptation to seasonality regulation in vertebrates (Dardente et al., 2014; Laudet, 2011; Sayre and Lechleiter, 2012). The extends to which these functions are shared across vertebrates remains unclear, but our data clearly suggest that the two first functions are active during clownfish metamorphosis: TH promotes the morphological and behavioral transformation of the larvae while alo promoting a metabolic shift in energy source.

We have studied in detail the action of TH on vision and metabolism and in a recent report, we uncovered their function for adult pigmentation pattern formation (Salis et al., 2021). Interestingly, our transcriptomic data obtained on whole larvae reveal that several other important biological processes such as bone mineralization and digestion are also changing during metamorphosis. This, together with the detailed analysis of vision and metabolism that we have performed, clearly illustrate the pleiotropic TH action during metamorphosis.

### A behavioral shift in vision controlled by TH

We observed a TH-induced shift in between short wavelength and long wavelength opsin gene expression at stage 4 during clownfish metamorphosis (Fig. 3). Interestingly, this shift occurs concomitantly with a behavioral change, namely a change in color preference. Larval fish prefers a blue background, in accordance with their natural pelagic environment, whereas juveniles prefer an orange background in accordance with the shallow, colorful and chromatically dynamic environment of the reef (Chiao et al., 2000). In addition, we showed that TH exposure decreases the preference for blue background on treated fish. In addition, the TH-treated larvae, adopted a benthic behavior characterized by low swimming activity close to the bottom of the experimental tank, which is in accordance with the benthic live style of anemonefish juveniles. Our data do not unequivocally demonstrate a direct effect of TH in color vision as many effects can also be elicited in neural circuit mechanisms for color vision (Baden, 2021). However, they strongly suggest that TH globally controls a visual shift that is ecologically critical for the recruitment success of juveniles.

Vision is one of the main sensory systems that has been studied by ecologists in the context of fish recruitment (Job and Shand, 2001; Shand, 1997). A shift in opsin gene expression, and in particular an increase of *opnlw* expression, has been observed in many fish species such as flatfishes, black bream or cichlids (Carleton et al., 2008; Ferraresso et al., 2013; Härer et al., 2017; Shand et al., 2008; Shao et al., 2017). However, up to date, there is only a partial understanding of the mechanistical control of this phenomenon and its integration with metamorphic transformation. In zebrafish, TH signaling has been shown to regulate the *opnlw* and mutation of the *TRβ* gene resulted in an absence of red cones but no link with an ecological transition was established (Suzuki et al., 2013). In surgeonfish, we have observed that shifts in retina structure and in the ability to visually recognize predators are both controlled by TH but no data on gene expression have been obtained so far, leaving the mechanism underpinning this recognition unclear (Besson et al., 2020).

The data we obtained here, are in accordance with what has been observed in mammals. In the mouse, (Eldred et al., 2018) have shown that TH controls the temporal switch between S and L/M cones which contains short and medium/long wavelength opsins, respectively. This suggests that the TH action on opsin gene expression and more generally visual function is a general phenomenon in vertebrates. By its ability to combine *in vitro* and field studies, the anemonefish model could allow to integrate the TH action on retina formation and more generally on sensory systems maturation in an ecologically relevant context (Roux et al., 2020).

### The metabolic transition is instrumental for metamorphosis completion

We observed a major metabolic transition during metamorphosis, and we provided evidence that TH is implicated in this transition. Namely, larvae preferentially use glycolysis and fermentation to produce energy whereas juveniles rely on fatty acid β-oxidation and citric acid cycle.

Our data show that there is a clear shift from glycolysis during early larval life (stage 1-3) to β- oxidation starting at stage 4 (Fig. 4). To put this finding in perspective we need to consider three parameters: (i) the changes in the organism growth; (ii) the transition of the fish into a new ecological life and (iii) the energy produced by each pathway. In glycolysis, one molecule of glucose produces a net excess of only 2 ATP, 2 NADH and 2 molecules of pyruvate while oxidation of one molecule of palmitic acid (C16:0) (one of the most common saturated fatty acid found in vertebrates), produces a net gain of 129 ATP molecules by β-oxidation (Wang et al., 2020). Therefore, utilizing fatty acid as a fuel source generates much more energy that sugar. During *A*. *ocellaris* post-embryonic development, we observe a massive increase in size soon after stage 4. Growth in vertebrates in highly demanding in energy and having the possibility for the larvae to utilize a fuel source that produces a large number of ATP is certainly an advantage. Moreover, after stage 4, the transforming fish must actively swim to find its juvenile habitat and once settled, will have to fight with congeners to be accepted in the colony. All this will require high energy demand, hence high ATP production explaining the transition to a more energy producing system.

In addition to this shift in energy production, we also reveal that TH also stimulates the production of complex lipids suggesting that the biochemical landscape of the juvenile is more elaborated than the larval one, because of TH action. These lipids can be used for fatty acid transport (acyl carnitine), energy storage (triacylglycerol), membrane constituents (ceramide, sphingomyelin, phosphatidyl ethanolamine) or signaling molecules (phosphatidylglycerol sphingomyelin) (Hulbert, 2021).

Interestingly, this metabolic transition is deeply linked to the morphological and behavioral changes that we describe in this paper. Indeed, when we modify lipid metabolism by using a LXR antagonist, we observed an acceleration of white bar appearance as well as a shift in opsin gene expression, two of the most salient endpoints of metamorphosis. This suggests that LXR is active during this period, in accordance with its expression peak at stage 4 and its up regulation by TH. LXR might be involved in slowing-down the process, ensuring that it is progressing concomitantly with the available energy production. In addition, by its known action in regulating elongases and desaturase genes, LXR is likely playing an important role in the complexification of fatty acids that we observed (Xie et al., 2021).

### The ecological function of thyroid hormone

Collectively our data suggest that TH is instrumental in integrating the complex remodeling occurring during metamorphosis with the available resources provided by the environment. This is highly relevant for a marine fish that similarly to a pelagic larvae, is living in a very dynamic environment characterized by patchy food resources (Seuront et al., 2001). It must be pointed out that the challenges faced by a tiny pelagic fish larvae are impressive: It must combine energy production from highly patchy food resources, mobilize its reserves in case of fasting and, at the same time, mature sensory systems that can detect future juvenile habitat, all this while avoiding predators! Our paper strongly suggests that TH plays a major role to orchestrate this complex transformation.

Interestingly, the fact that metabolism must be coupled with developmental transition to fulfill the energy requirements during organisms’ life cycle, have been previously pointed out in the context of insect metamorphosis. (Nishimura, 2020) have revealed that the programed regulation of metabolism by steroid hormones control the key steps during drosophila metamorphosis. This is remarkably similar to the effects we observed here with the major difference that, in contrast to insect pupa, the transforming fish larva rely on an external food source. However, in both cases, the coordination of the larval transformation must integrate both environmental and internal conditions. Adjusting metabolic regulation is a major tool for achieving this.

Our data are also relevant to consider in the light of previous work done on stickleback. These fish can live in either marine or freshwater environment and the energy resource available in these two situations are drastically different. Interestingly, Kitano et al., 2010 have shown that, in freshwater sticklebacks there was a fine tuning of numerous physiological and metabolic traits that ensure a reduced energy expenditure. These changes are driven by TH, as a genetic variant expressing low level of TSH, the main hypothalamic peptide controlling TH production, has been fixed in these populations (Kitano et al., 2010). In these freshwater sticklebacks in which TH level is low, there is also evidence of a shift in opsin gene expression linked to local adaptation to different light environment (Rennison et al., 2016). This suggests that, in stickleback as in anemonefish, there is a tight integration of metabolic and sensory changes, and that TH may be instrumental in this process. This case provides a genuinely nice illustration of a still unappreciated potential for TH regulation to induce pleiotropic changes, allowing a multi-level regulation of a developmental transition (Laudet, 2010).

Our model has also a general significance regarding the effect of climate change and pollution in animal populations. Indeed, the sensitivity of metamorphosis to environmental stressors has been recently emphasized (Lowe et al., 2021). Metamorphosis being a life-history transition with abrupt oncogenic changes tightly connecting to environmental conditions can make young life stages vulnerable to stressors. Indeed, we demonstrated previously in convict surgeonfish, that temperature and pollution both synergistically disrupt TH signaling affecting the ability of the young juveniles to perform their ecological function (Besson et al., 2020). Our study therefore suggests that more research is needed to decipher how stressors affects not only metamorphosis endpoints but also the metabolic regulations that ensure its correct progression.

## Material and Methods

### *Amphiprion ocellaris* maintenance and rearing

*Amphiprion ocellaris* larvae were obtained from breeding paired maintained as described in Roux et al. (2021) in a rearing structure. Reproductive pairs laid eggs every two weeks allowing us to rear larvae regularly for the purpose of this study.

### Transcriptomic analysis

#### RNA extraction

Three larvae per stage are sampled making a total of 21 samples. Larvae are euthanized in a MS222 solution (200 mg/l), photographed for stage identification, and kept in RNAlater prior to RNA extraction. Total RNA is extracted from whole individual larval body using a Maxwell®16 System (Promega, Madison, USA) and following the manufacturer’s instructions. RNA integrity and concentration is verified with an Agilent 2100 Bioanalyzer (Agilent Technologies, Santa Clara, USA) and only samples with RIN values equal to, or above 8 are used. In our case, all samples had RIN values above 9.

### RNA-Seq libraries preparation and sequencing

RNA-Seq libraries and sequencing are performed by the IGBMC in Strasbourg on an Illumina HiSeq4000 sequencer using a stranded protocol with paired-end 2x100bp.

### Software

Transcriptomic analyses are performed by the Altrabio Company (Lyon, France) using the following software: FasQCversion 0.11.8 and fastq_screen version 0.13.0, Salmon version 0.14.1, R version 3.6.1 (2019-07-05), x86_64-apple-darwin15.6.0.

### Pre-treatment (read quality, quantification, filtering, normalization)

Quality of raw reads is assessed using the FastQC quality control tool1. Sample contamination is assessed using the fastq_screen quality control tool2. The Bowtie2 indexes for Amphiprion ocellaris are generated from Ensembl “Amphiprion_ocellaris.AmpOce1.0.dna.toplevel.fa” file. Amphiprion ocellaris transcript sequences and annotations (Ensembl release 97) are downloaded from the Ensembl website3 (files Amphiprion_ocellaris.AmpOce1.0.cdna.all.fa, Amphiprion_ocellaris.AmpOce1.0.ncrna.fa and Amphiprion_ocellaris.AmpOce1.0.97.gtf). Some genes which had no official symbols in Ensembl are given the following symbols: ENSAOCG00000023317 =tg, ENSAOCG00000008531 = mct8, ENSAOCG00000005709 = dio3a, ENSAOCG00000017526 = edn3b, and ENSAOCG00000019494 = bnc2. Transcript expression quantification is performed from the raw read data (FASTQ files) using Salmon4 (with parameters --gcBias, --seqBias, --validateMappings, -- discardOrphansQuasi, and --consistentHits). They are then summarized as gene counts using function tximport from package tximport5 (with parameters type=“salmon” and countsFromAbundance=“lengthScaledTPM”). Genes that did not have more than 0.502 count per million counts in at least three samples are filtered out. To generate the normalized signals, the effective library sizes are first computed using function estimateSizeFactors from package DESeq26. The raw count values are then transformed using a variance stabilizing transformation (function VarianceStabilizingTransformation from package DESeq2 with parameter blind=TRUE).

### Hierarchical clustering

Hierarchical clusterings of samples are performed using the Wards agglomerative method, passing the euclidean distances between samples to function hclust from package stats (with parameter method=“ward.D2”). Cluster stability is estimated by multiscale bootstrap resampling using function pvclust from package pvclust7 (with parameter nboot=10000). Hierarchical clusterings of genes are performed using the complete agglomerative method, passing the Euclidean distances between centered (for expression data) or uncentered (for fold change data) and scaled gene signals with function hclust from package stats.

### Principal component analysis

Principal Component Analyses (PCA) of sample expression levels are performed with gene signals centered but not scaled (using function prcomp from package stats). When displayed, gene coordinates correspond to these genes’ correlations with the presented components. Only the genes that contribute most to the components are displayed.

### TH dosage

To determine at which stage TH are surging and marking the beginning of metamorphosis, pools of five clownfish larvae are sampled in triplicate at each of the seven developmental stages identified in Roux et al. (2019). THs are extracted from dry-frozen larvae (previously euthanized in a 200 mg/l solution of MS-222) following the protocol developed by Holzer et al. (2017) and adapted from previous TH extractions in teleost fishes (Einarsdóttir et al., 2006b; Kawakami et al., 2008; Tagawa and Hirano, 1989). TH concentrations are measured by a medical laboratory in Perpignan (Médipole) using an ELISA kit (Access Free T3, T4, Beckman Coulter).

### *In situ* hybridization

Digoxigenin RNA probes were synthesized using the T3/T7 Transcription Kit (Roche; Supp. Table 1). Larvae were collected, euthanized in MS222 at 200 mg/L and fixed 12 hours in 4%. paraformaldehyde diluted in PBS (phosphate-buffered saline). Samples were subsequently dehydrated stepwise in PBS/ethanol, and then put three times 10 min in butanol 100% and finally in two bath of paraffin (respectively 1 and 4 hours) before being embedded in block. Embedded larvae were sectioned transversally at 7µm using Leica Biosystems RM2245 Microtome the day before starting ISH. The samples were then treated as in Thisse, Thisse, Schilling, and Postlethwait (1993).

### Behavioral test for visual perception

Choice experiments are conducted on stage 2 and stage 5 to determine if there is a shift in visual perception before and after metamorphosis. Thus, a dual choice aquarium (measuring 25x10x10 cm) is built with white opaque comassel, fitted with transparent plexiglass on the sides (Fig. 3C). The aquarium is divided into three equal compartments (Fig. 3C). The larvae are given the choice between a short wavelength color: blue and a long wavelength color: orange. Each color is placed close to the plexiglass sides. Behavioral experiments are conducted in a dark room where the choice chamber is installed under a light ramp to ensure homogenous distribution of the light over the device. All the larvae are tested individually. The larvae are introduced carefully in the choice chamber without the colored panels and left for acclimation for 2 min. They are free to explore the three compartments. After acclimation and once the larvae are localized in the central compartment, blue and orange panels are installed, and the time spent by the larvae in each compartment is recorded during a 5 min period. We established that a larva preferred a color when it is spending most of its time in the compartment of the given color (Fig. 3C). Color panels are inverted after each tested larva to ensure that the choice is due to the color and not to a defect in the experimental device. Results are expressed in the mean percentage of time spent in each compartment. A Kruskall- Wallis test is performed to determine if the difference in the amount of time spent between larvae (stage 2 vs stage 5, and DMSO vs T3) is significantly different in each compartment.

### T3 treatments

To test the effects of THs on the metamorphosis of clownfish A. ocellaris, larvae were exposed to various concentration of T3 using the low-rearing volume protocol developed by Roux et al., (2021). Briefly, larvae were exposed to three different concentrations of a mix of T3 and iopanoic acid (IOP) to induce metamorphosis. IOP was added to block the action of the deiodinases and avoid the degradation of the added T3 (Holzer et al., 2017). Larvae were exposed to increasing doses of T3: 10^-8^M, 10^-7^M, 10^-6^M (the IOP concentration remained constant at 10-7M), and to DMSO (as control) diluted at 1:1000. Larvae were sampled at 12h, 24h, 48h and 72h for gene expression analysis using nCounter technology from Nanostring. Exposition started at stage 3 before the beginning of the metamorphosis and lasted for 12h, 24h, 48h and 72h. Larvae were introduced by groups of 10 in 800 ml beakers placed in a water bath to maintain the temperature at 27°C. Larvae were fed 3 times a day with rotifers at a final concentration of 10 rotifers/ml and once a day with nauplii of artemia (Roux et al., 2021). The algae *N. oculata* was added (500µl/beaker) at each feeding to create a green environment and improve survival rates. During the experiment, water changes of 100 ml (with addition of each treatment to maintain constant concentrations) were done daily to ensure water quality. Larvae were sampled after 12h, 24h, 48h, and 72h of exposition, euthanized in a MS222 solution (200mg/l) and photographed under a Zeiss stereomicroscope (V20 discovery Plan S) equipped with an Axiocam 105 camera for morphological analysis. Three replicates composed of pools of two larvae were then kept in RNA later at -20°C for gene analysis using nCounter technology from Nanostring.

### Gene expression measurements using nCounter technology Probe synthesis

Probes of 100 nucleotides were designed by Integrated DNA Technology for all the genes investigated in this study (Supp. Table 2). However, no probes could be designed that specifically targeted *TRαa, TRαb*, *rh2a-1* and *rh2a-2* as sequences were too similar. For this reason, only one pair of probes has been designed and targeting both *TRα* and *rh2a* genes. Seven reference genes were chosen for normalization and expression analysis: *Tuba1, PolD2, g6pd, eif4a3, tbp, rpl7* and *rpl32*.

### Sample processing

Samples of each experiment were processed in multiplexed reaction including, in each case, six negative probes (to determine the background) and six positive control probes. Hybridization reactions lasted 16 hours at 67°C. Data were then imported into the nSolver analysis software (version2.5) for quality checking and normalization of data according to NanoString analysis guidelines, using positive probes and 7 housekeeping genes (*Tuba1, PolD2, g6pd, eif4a3, tbp, rpl7* and *rpl32*) (Kulkarni, 2011).

### Statistical analysis

After normalization, analysis of the difference between treatments for each experiment was conducted using ANOVA (R software 3.2.3 version) and fold change calculated on a log2 scale. As multiple comparison was used, p-value was adjusted using the Benjamini & Hochberg method (Ferreira and Zwinderman, 2006).

### Lipid composition measurements and analysis

Larvae were sampled at stage 3, 4, 5 and 6 and starved for one hour before being flash frozen in liquid nitrogen and stored at -80 C° until analysis. Three individuals were collected for each condition and lipid extraction was done on each individual.

Samples were analyzed as previously described (Weir et al., 2013), briefly, samples were thawed and lyophilized to remove all liquid. Samples were then reconstituted prior to extraction in 10 μl of water and 10 μl of internal standard mix (ISTD) was added to each sample. Lipids were extracted by adding 200 μl chloroform/methanol (2:1) and sonicated in a bath for 30 minutes before the supernatant was transferred to a 96-well plate and dried under vacuum in a SpeedVac Concentrator (Savant). Samples were then reconstituted with 50 μl water saturated butanol and 50 μl methanol with 10 mM ammonium formate and analysed by LC ESI-MS/MS using an Agilent 1200 LC system and an ABSciex 4000 Qtrap mass spectrometer. Data was processed using MultiQuant 2.1 software. Lipids are presented as pmol/mg protein. The lipid standards used in this analysis were as previously described (Weir et al., 2013).

Statistical analysis on developmental stage was performed using one way ANOVA with R software for each lipid followed by a multiple pairwise comparison performed by pairwise t-test in case of significant differences (p-value<0.05) (Fraher et al., 2016).

Similar analysis was performed on larvae treated for 72hours at 10^-8^ M and 10^-7^ M T3(mixed with 10^-7^ M IOP). Sampling was performed as explained previously. Statistical analyses were performed on comparing each T3 concentration with DMSO control condition using the test of Student with R software.

### LXR experiments

Specificity of a human LXR antagonist SR9243 was tested in reporter cell lines expressing the ligand binding domains of zebrafish receptors LXR, PPARγ and PXR. Briefly, SR9243 was tested alone for its agonistic activity in HG5LN-zfLXR, -zfPPARγ and zf-PXR cells (Creusot et al., 2020; Garoche et al., 2021). For its antagonistic activity on -zfLXR, -zfPPARγ and zf-PXR cells, SR9243 was tested in combination with T091317 30 nM, GW3965 3 μM and clotrimazole 0.1 μM respectively (Supp. Fig. 10).

Larvae of *A. ocellaris* were sampled at stage 3 and treated for 5 days with DMSO (control condition) and LXR antagonist SR9243 at 10^-7^ M. Experiment was conducted following the protocol described in Roux et al., 2021 and detailed previously. Gene expressions was obtained by RT-qPCR (PrimeScript transcriptase, Takara, SYBRgreen) and normalized with Pfaffle equation using two housekeeping genes (*rpl7, rpl32*) (Stahlberg et al., 2004). Specific primers were designed using Primer3 software (Untergasser et al., 2012). Primers sequenced are summarized in Supp. Table 1. Significant difference of gene expression level between DMSO control and SR9243 conditions was assessed by using the test of Student with R software. Difference between the number of individuals displaying white bars in both conditions was assessed by a Chi² test performed with R software.

### Amphiprion percula RNA sequencing

*Amphiprion percula* new recruit were sampled in Kimbe, Papua New Guinea in both *Heteractis magnifica* and *Stichodactyla gigantea.* A total of three new recruits per sea anemone were euthanized in MS222 solution and stored in RNAlater until RNA extraction and RNA sequencing of the whole fish individually (Salis et al. 2021). Potential adapter contaminations were removed and trimmed to obtain raw reads with cutadapt (v.1.13; (Martin, 2011)) and sickle (v1.29, (Joshi and Fass, 2011)), respectively. The processed reads were mapped against *A. percula* reference genome (Ensembl ID: GCA_003047355.1; (Lehmann et al., 2019)) using HiSat2 (v.2.1.0; (Kim et al., 2015)). Raw counts for each gene were obtained with HTSeq (htseq-count, v.0.9.1; (Anders et al., 2015)), using the available gene annotation of the *A. percula* reference genome (Salis et al. 2021). Raw counts were then normalized as transcript per million to assess the difference in expression levels for gene involved in vision and metabolic processes using the test of student. Principal component analysis was also performed on each biological process separately.

## Aknowledgements

We thank the Service Mutualisé d’Aquariologie of the Observatoire océanologique de Banyuls-sur-mer who take care of the clownfish husbandry. We thank the Bio2mar platform for their expertise for RNA extraction and RNA integrity analysis. NCounter analysis were performed by Nathalie Saint Laurent, Carine Valle, and Marie Tosolini from the CRCT of Toulouse and lipid analysis were performed by Oswald Quehenberger, Aaron Armando from the Lipidomics Core service core (University of California, San Diego). We also thank Serge Planes for the sampling of *A. percula* recruit in Papua New Guinea. We thank Roger Huerlimann, Marleen Klann and Tim Ravasi for critical reading of the manuscript.

## Author contribution

NR and VL wrote the manuscript with contributions from YG, KG, DL and LB. NR, VL, DL, LB designed the whole study with the help of YG and KG for the metabolic aspects. Transcriptome assembly was performed by SB and analysis were performed by SB and NR. NR performed all the pharmacological treatments with the help of SM, MD, YT and FL. NR performed gene expression and behavioral experiments linked to visual perception and metabolism (with the help of SM and YT). YG and KG brought their expertise on metabolic regulation and lipid metabolism. YG helped in the lipid analysis. NR, SM and YT conducted experiments on lipid metabolism gene expression. PS generated the transcriptomic data set of *Amphiprion percula* recruits sampled in Kimbe, Papua New Guinea. ABa and NR analyzed *A. percula* transcriptome. NR and MR conducted thyroid hormone measurments. Abo performed LXR antagonist activity experiment on zebrafish.

## Ethics Approval

All experiments conducted in this study were done under the approval from the C2EA-36 Ethics Committee for Animal Experiment Languedoc-Roussillon (CEEA-LR - approval N°A6601601) as well as following the Animal Experiment Regulations at Okinawa Institute of Science and Technology Graduate University (approval N°20052605).

## Supplementary Figure legends

**Supplementary Figure 1:**
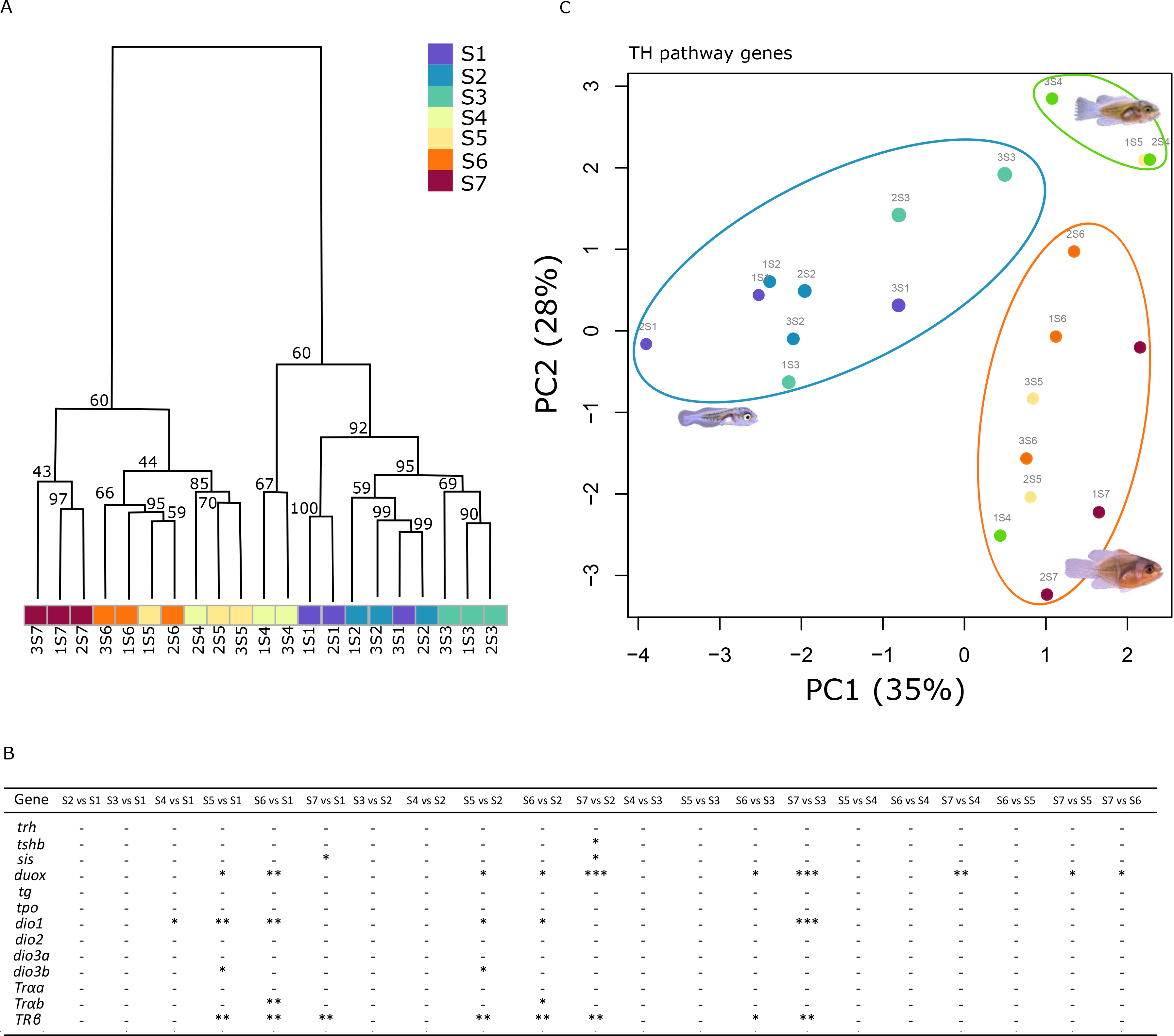
(A) Principal component analysis generated using TH signaling pathway gene expression (obtained by RNA sequencing, n=3 per stage) showing a clear separation before and after metamorphosis. (B) Table summarizing the TH signaling genes significantly differentially expressed between each stage.

**Supplementary Figure 2:**
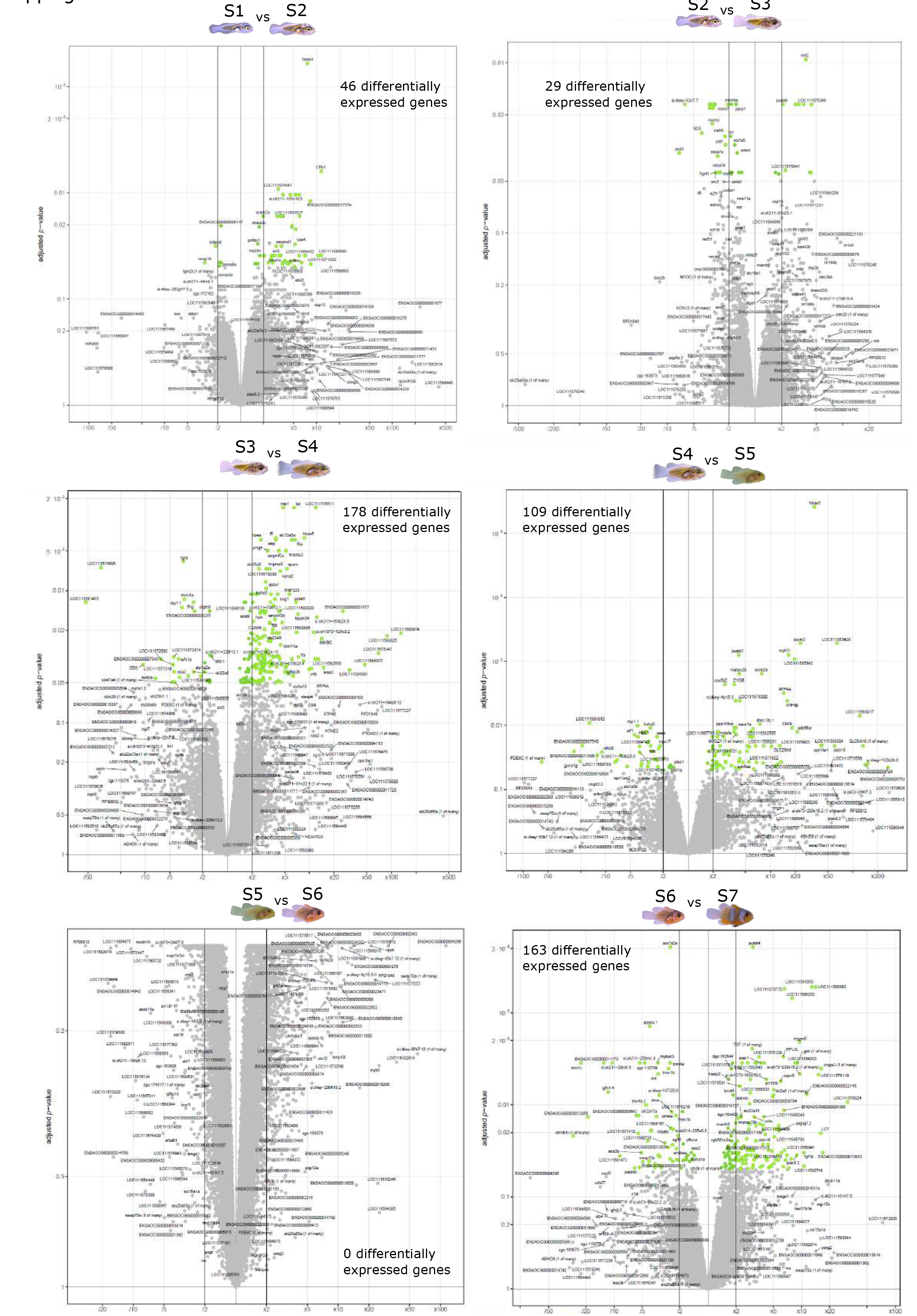
Volcano plots showing the number of gene significantly differentially expressed between each contiguous developmental stage (S1: stage 1; S2: stage 2; S3: stage 3; S4: stage 4; S5: Stage 5; S6: stage 6 and S7: stage 7).

**Supplementary Figure 3:**
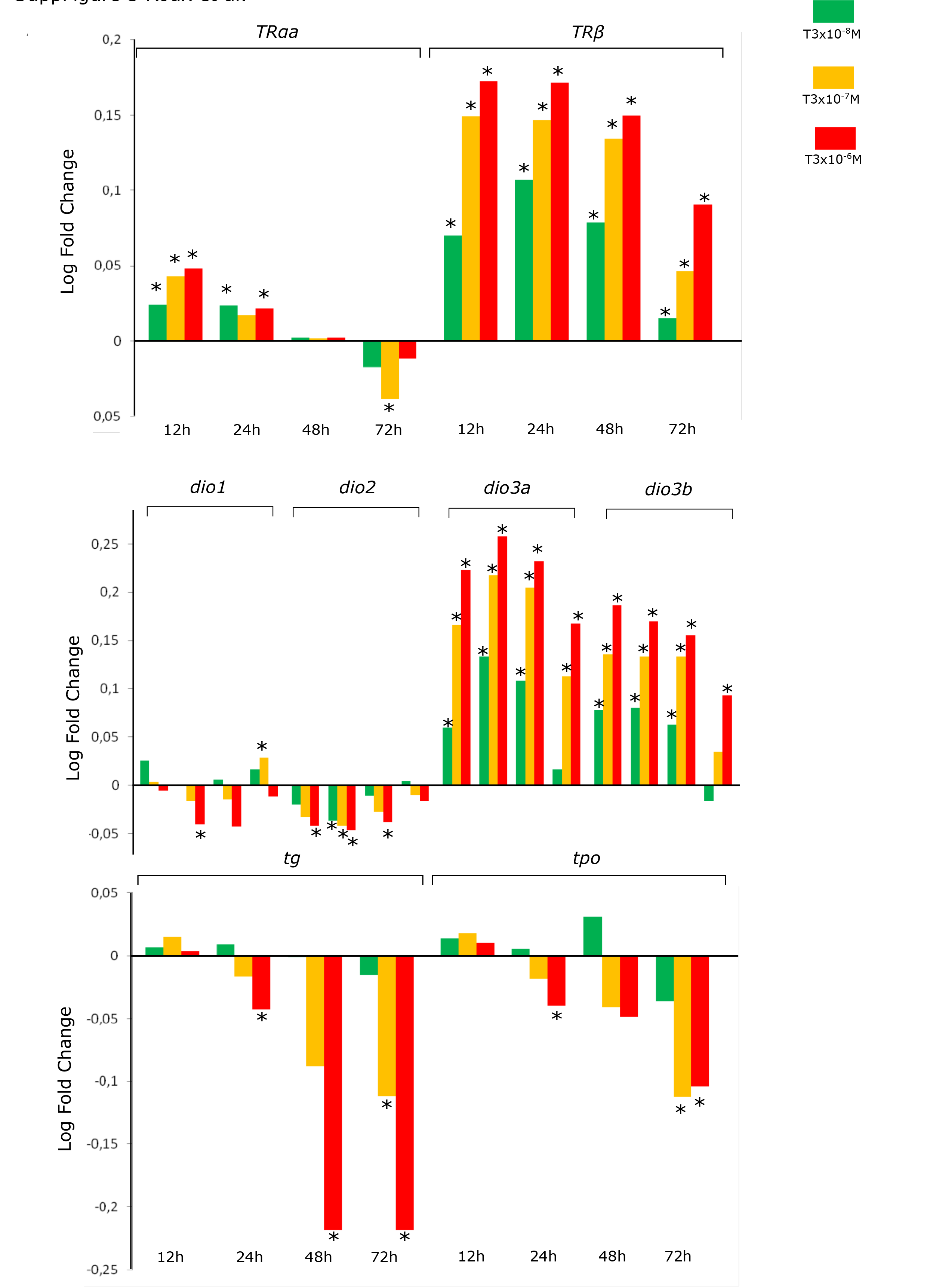
Graphics showing the effect of thyroid hormone (T3 at 10^-8^, 10^-7^, 10^-6^ M) after 12, 24, 48 and 72hours of treatment on the expression levels (fold change) for genes involved in (A) thyroid hormone receptors (*TRa, TRβ),* (B) genes involved in TH metabolism (*dio1, dio2, dio3a, dio3b*) and (C) TH synthesis (*tg, tpo,* sis),

**Supplementary Figure 4:**
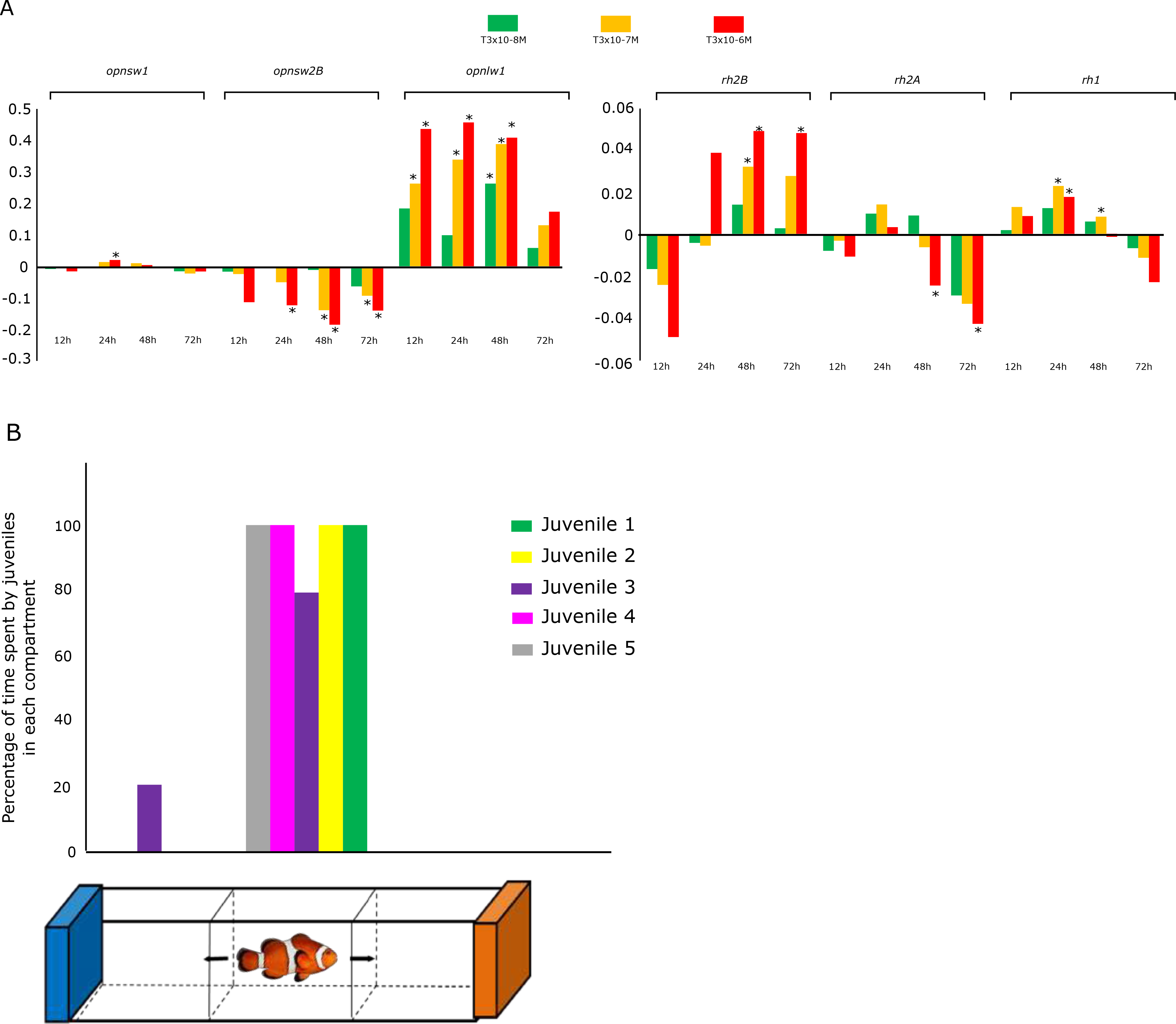
(A) Graphics showing the expression levels of *opnlw1, opnsw1, opnsw2B, Rh1, Rh2A* and *Rh2B* when larvae were treated for 48 hours to increasing concentrations of T3 (10^-8^, 10^-7^, 10^-6^ M). Stars (*) Indicate the significant differences between treatment and control DMSO. (B) Graphic showing the percentage of time spent by five 30 days post hatching (dph) clownfish juveniles in the dual choice chamber when given the choice between orange and blue colors.

**Supplementary Figure 5:**
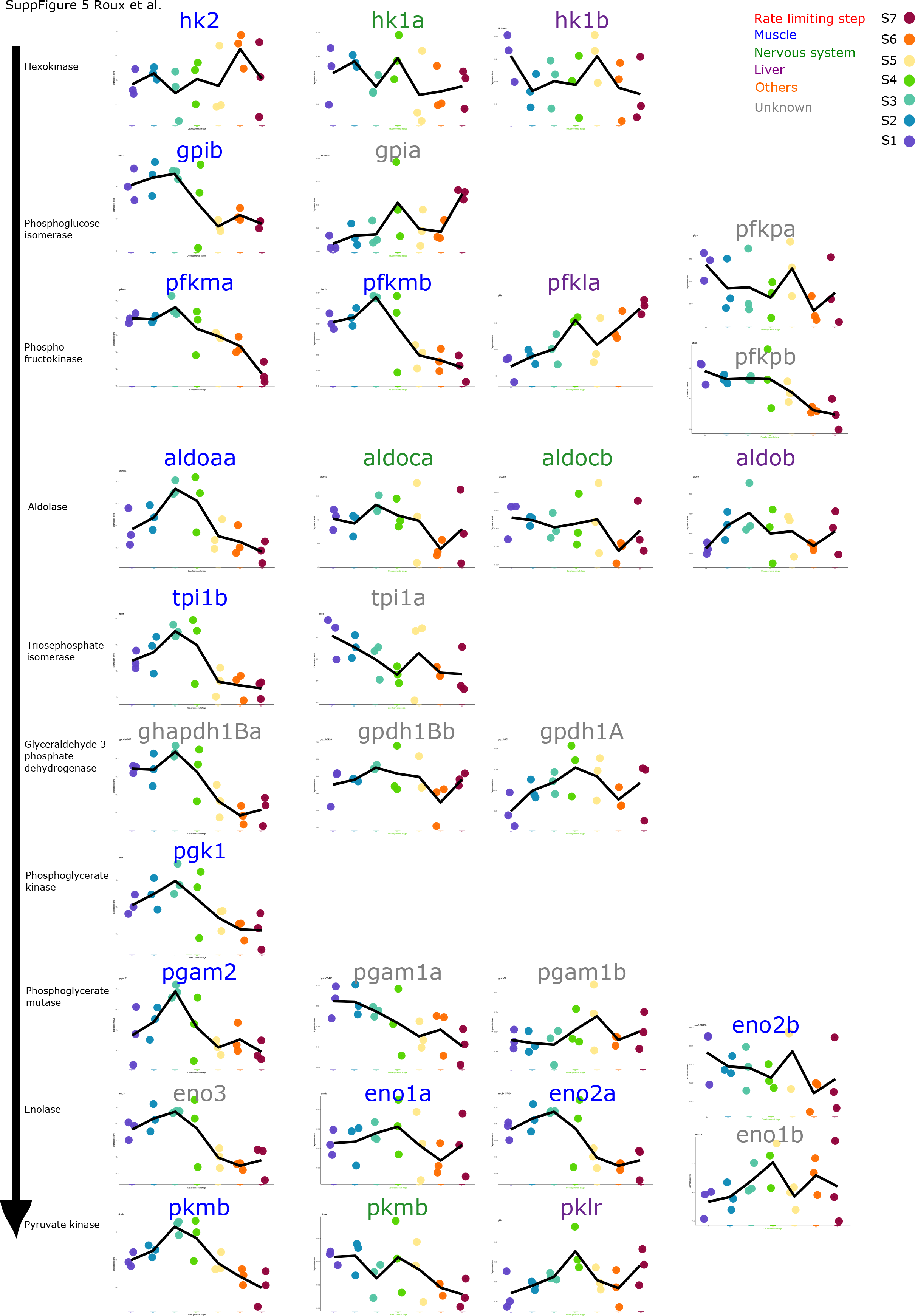
Figure showing the expression levels of all the genes encoding for the enzyme involved in glycolysis. Each color indicates the organs in which these genes are known to be preferentially expressed in zebrafish. (S1: stage 1; S2: stage 2; S3: stage 3; S4: stage 4; S5: Stage 5; S6: stage 6 and S7: stage 7).

**Supplementary Figure 6:**
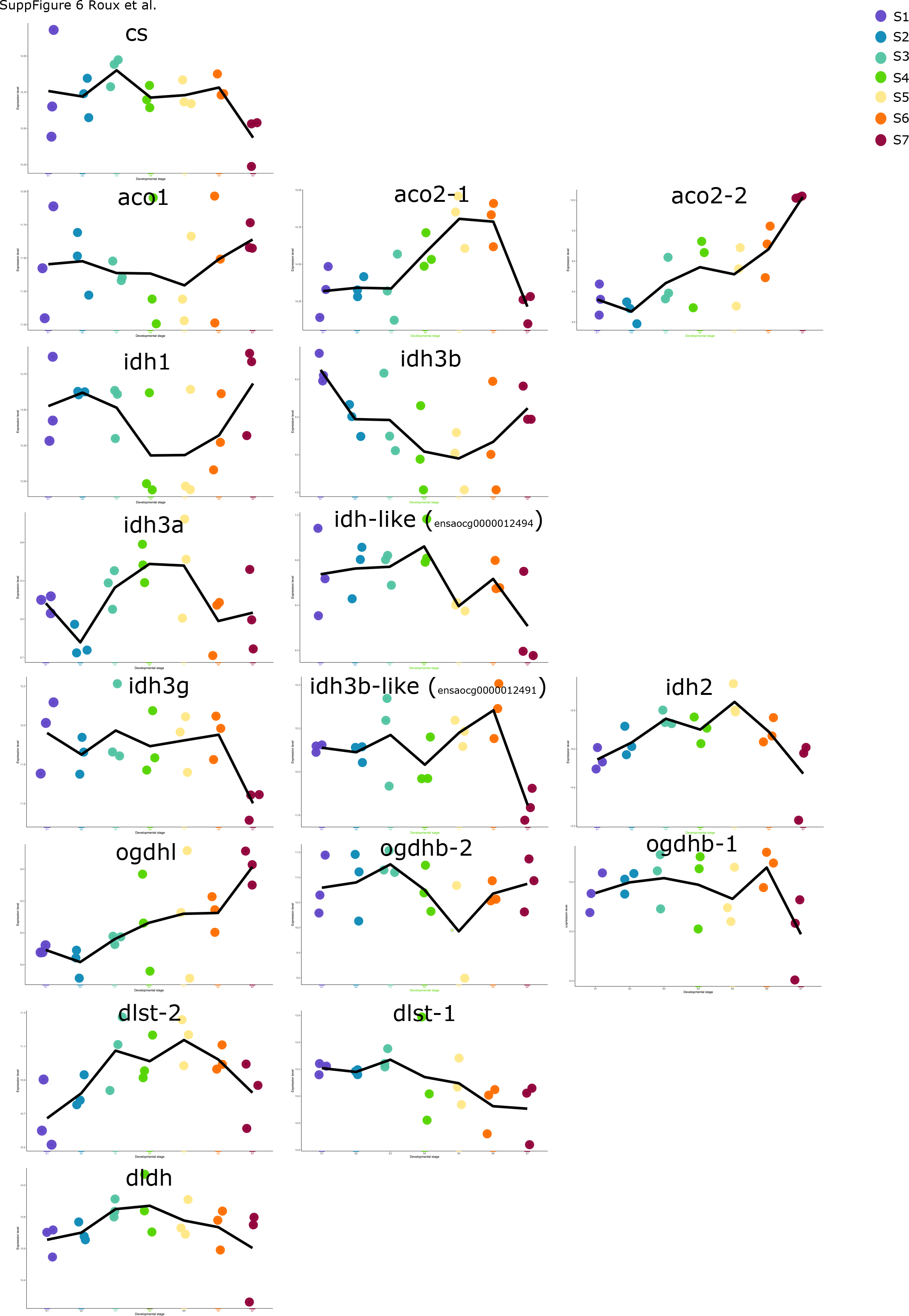
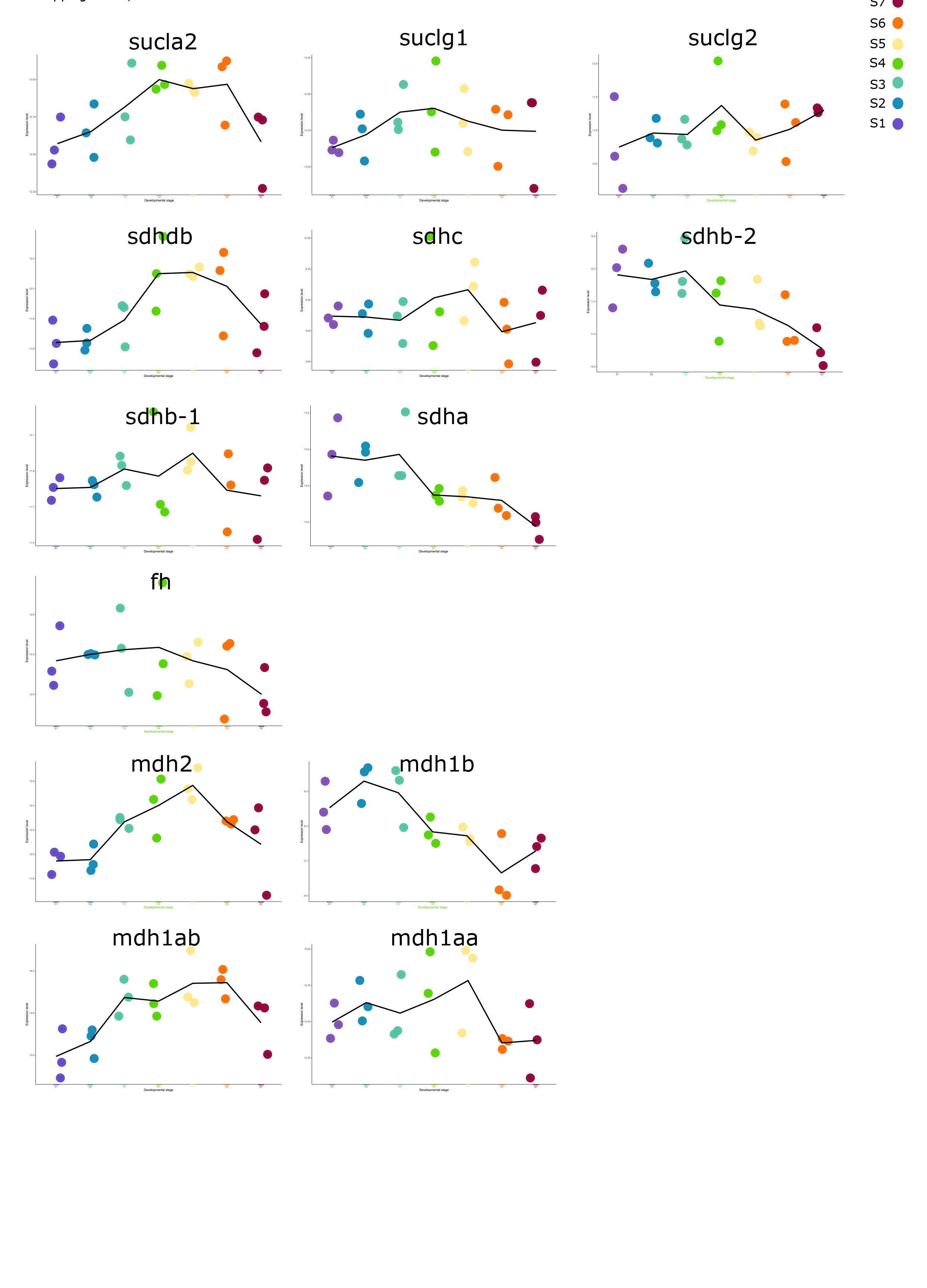
Figure showing the expression levels of all the genes encoding for the enzyme involved in citric acid cycle during *A. ocellaris* development (S1: stage 1; S2: stage 2; S3: stage 3; S4: stage 4; S5: Stage 5; S6: stage 6 and S7: stage 7).

**Supplementary Figure 7:**
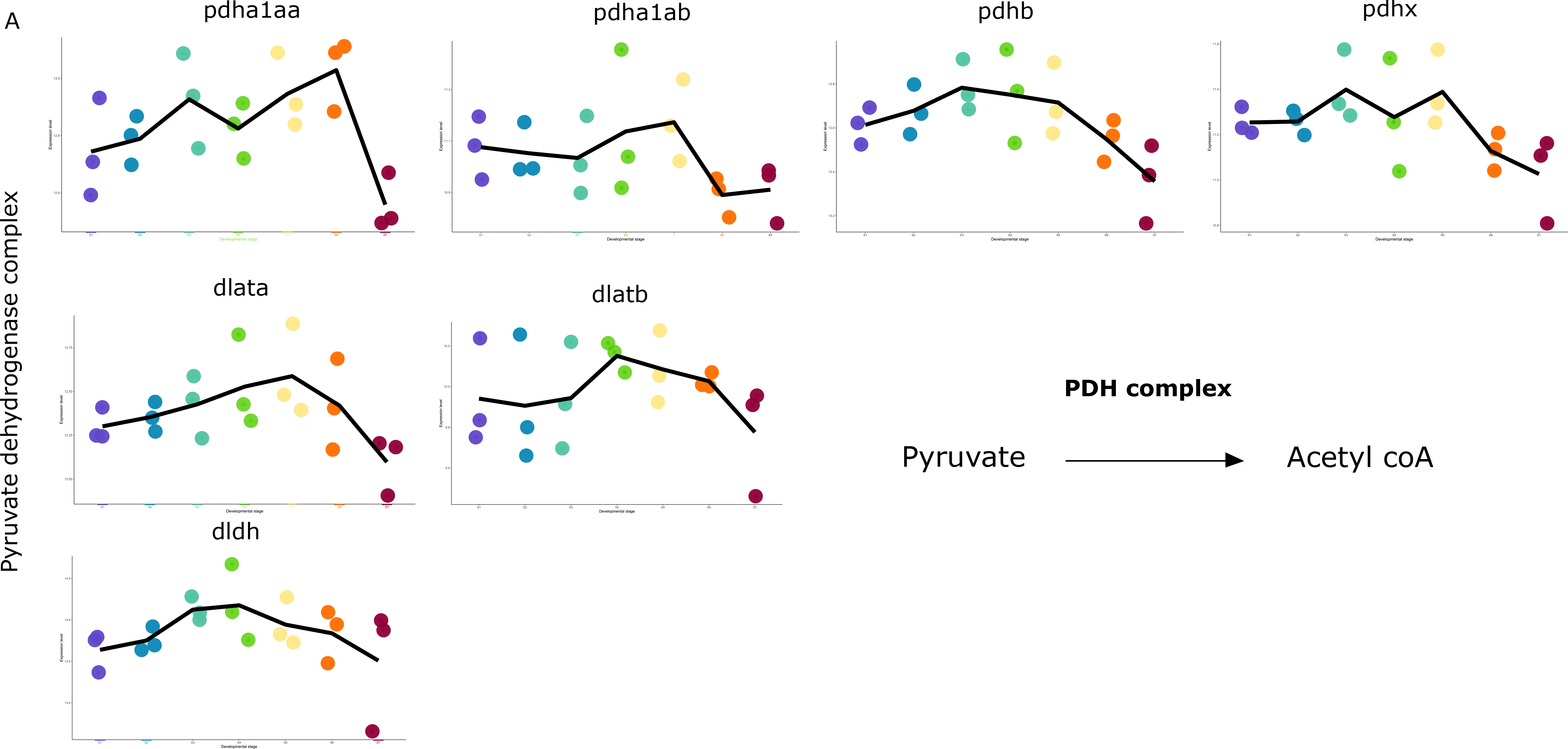
(A) the expression levels (obtained by RNA sequencing) during larval development of genes encoding for enzymes of the pyruvate dehydrogenase complex (catalyzing the transformation of pyruvate in acetyl coA).

**Supplementary Figure 8:**
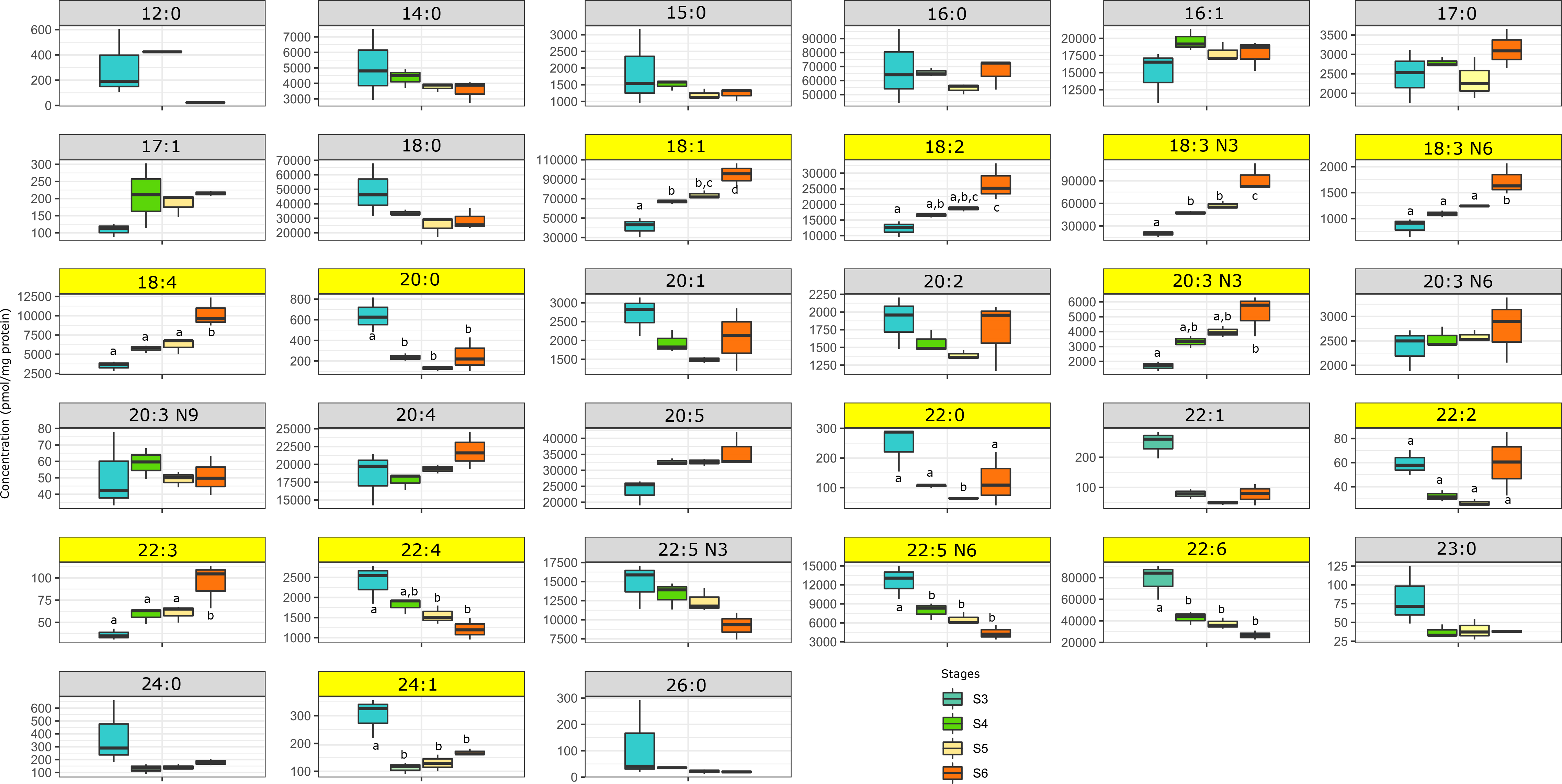
Fatty acid concentration measured at stage 3, 4, 5 and 6 (n=3 per condition). Fatty acid with concentration significantly different between stages are highlighted in yellow and different letters indicate significant differences between stages.

**Supplementary Figure 9:**
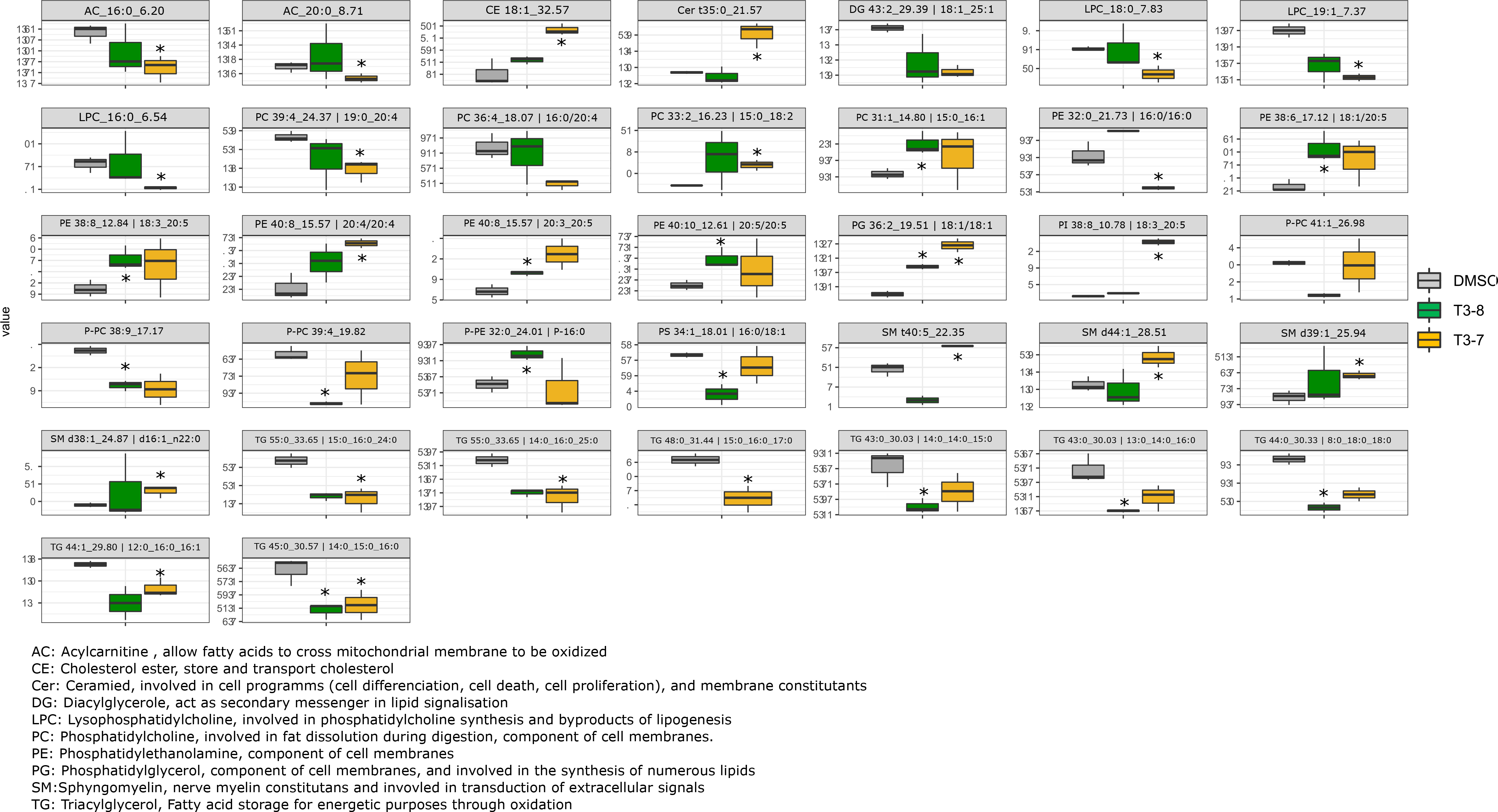
Complex fatty acid concentration whose differences are significantly different (*) between DMSO control larvae and T3 treated larvae (10^-8^, 10^-7^M, n=3 per condition).

**Supplementary Figure 10:**
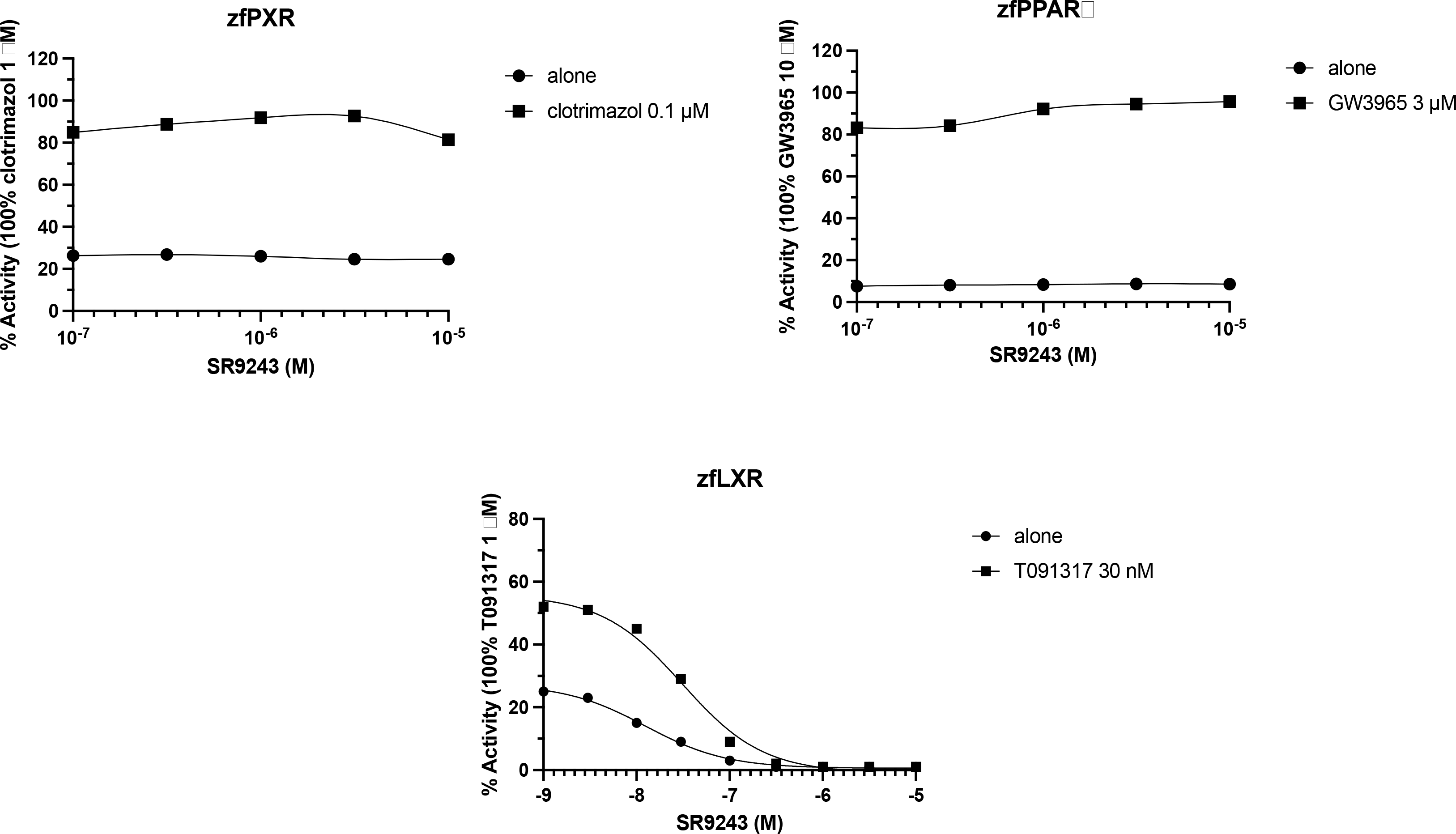
Antagonist Activities of zebrafish nuclear receptors (PXR, PPARg, LXR) against SR9243 confirming that SR9243 is a specific LXR antagonist.

**Supplementary Table 1:**
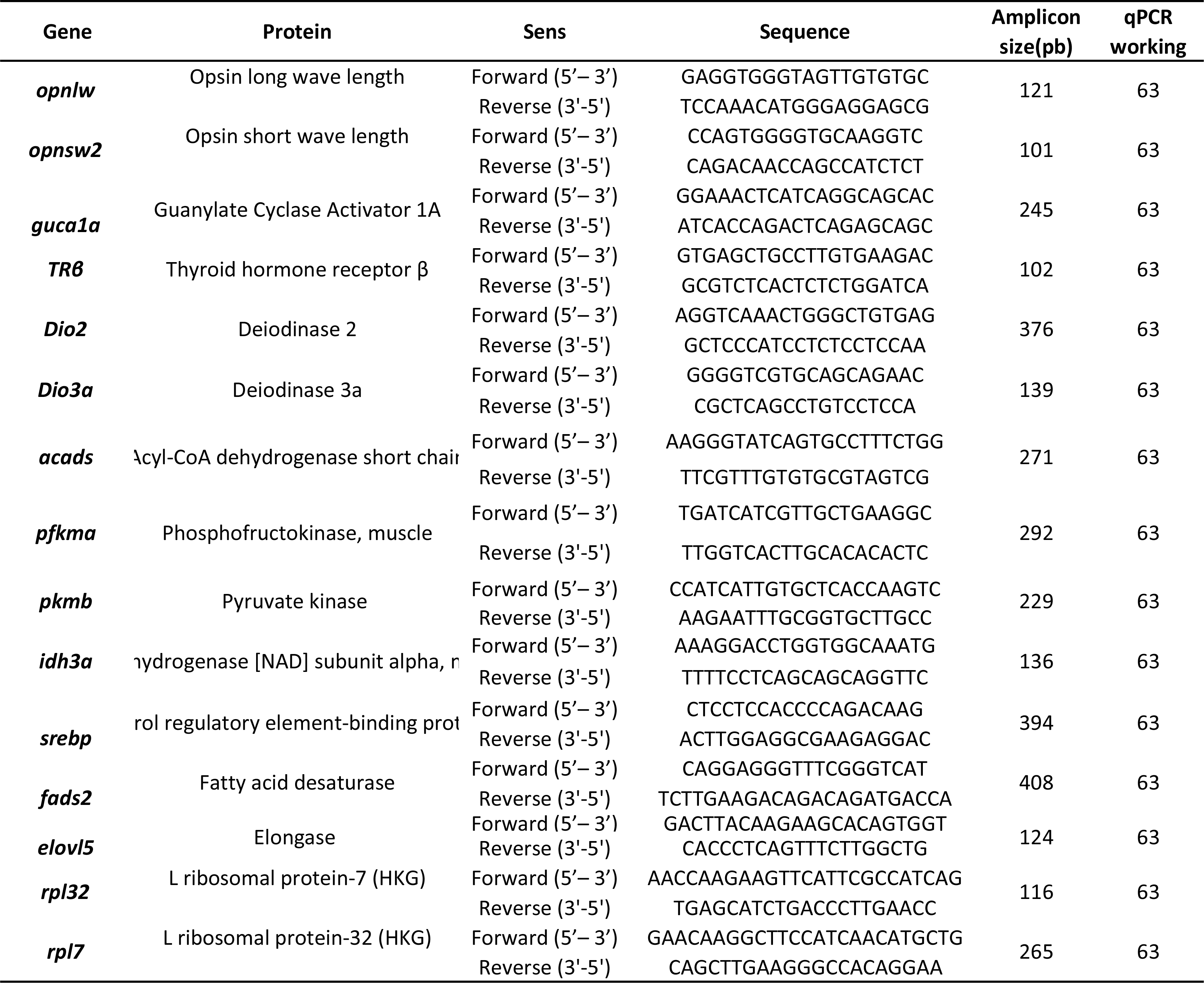
Primers sequence for RT-qPCR analysis of the LXR antagonist experiment.

## References

Alves Martins, D., Rocha, F., Martínez-Rodríguez, G., Bell, G., Morais, S., Castanheira, F., Bandarra, N., Coutinho, J., Yúfera, M., and Conceição, L.E.C. (2012). Teleost fish larvae adapt to dietary arachidonic acid supply through modulation of the expression of lipid metabolism and stress response genes. Br J Nutr 108, 864–874.

Anders, S., Pyl, P.T., and Huber, W. (2015). HTSeq--a Python framework to work with high-throughput sequencing data. Bioinformatics 31, 166–169.

Baden, T. (2021). Circuit mechanisms for colour vision in zebrafish. Current Biology 31, R807–R820.

Barth, P., Berenshtein, I., Besson, M., Roux, N., Parmentier, E., Banaigs, B., and Lecchini, D. (2015). From the ocean to a reef habitat: how do the larvae of coral reef fishes find their way home. VIE ET MILIEU-LIFE AND ENVIRONMENT 95, 91–100.

Besson, M., Feeney, W.E., Moniz, I., François, L., Brooker, R.M., Holzer, G., Metian, M., Roux, N., Laudet, V., and Lecchini, D. (2020). Anthropogenic stressors impact fish sensory development and survival via thyroid disruption. Nat Commun 11, 3614.

Bianco, A.C., and Kim, B.W. (2006). Deiodinases: implications of the local control of thyroid hormone action. J Clin Invest 116, 2571–2579.

Buchholz, D.R. (2015). More similar than you think: Frog metamorphosis as a model of human perinatal endocrinology. Dev Biol 408, 188–195.

Buston, P. (2003). Social hierarchies: size and growth modification in clownfish. Nature 424, 145–146.

Campinho, M.A. (2019). Teleost Metamorphosis: The Role of Thyroid Hormone. Frontiers in Endocrinology 10, 383.

Carleton, K.L., Spady, T.C., Streelman, J.T., Kidd, M.R., McFarland, W.N., and Loew, E.R. (2008). Visual sensitivities tuned by heterochronic shifts in opsin gene expression. BMC Biol 6, 22.

Chang, J., Wang, M., Gui, W., Zhao, Y., Yu, L., and Zhu, G. (2012). Changes in thyroid hormone levels during zebrafish development. Zoolog Sci 29, 181–184.

Charalambous, M., and Hernandez, A. (2013). Genomic imprinting of the type 3 thyroid hormone deiodinase gene: Regulation and developmental implications. Biochimica et Biophysica Acta (BBA) - General Subjects 1830, 3946–3955.

Chiao, C.-C., Cronin, T.W., and Osorio, D. (2000). Color signals in natural scenes: characteristics of reflectance spectra and effects of natural illuminants. J. Opt. Soc. Am. A, JOSAA 17, 218–224.

Cortesi, F., Musilová, Z., Stieb, S.M., Hart, N.S., Siebeck, U.E., Cheney, K.L., Salzburger, W., and Marshall, N.J. (2016). From crypsis to mimicry: changes in colour and the configuration of the visual system during ontogenetic habitat transitions in a coral reef fish. J Exp Biol 219, 2545–2558.

Creusot, N., Gassiot, M., Alaterre, E., Chiavarina, B., Grimaldi, M., Boulahtouf, A., Toporova, L., Gerbal- Chaloin, S., Daujat-Chavanieu, M., Matheux, A., et al. (2020). The Anti-Cancer Drug Dabrafenib Is a Potent Activator of the Human Pregnane X Receptor. Cells 9.

Dardente, H., Hazlerigg, D., and Ebling, F. (2014). Thyroid Hormone and Seasonal Rhythmicity. Frontiers in Endocrinology 5.

Darias, M.J., Zambonino-Infante, J.L., Hugot, K., Cahu, C.L., and Mazurais, D. (2008). Gene expression patterns during the larval development of European sea bass (Dicentrarchus labrax) by microarray analysis. Marine Biotechnology 10, 416–428.

Darras, V.M., and Van Herck, S.L.J. (2012). Iodothyronine deiodinase structure and function: from ascidians to humans. J Endocrinol 215, 189–206.

Deal, C.K., and Volkoff, H. (2020). The Role of the Thyroid Axis in Fish. Frontiers in Endocrinology 11, 861.

Einarsdóttir, I.E., Silva, N., Power, D.M., Smáradóttir, H., and Björnsson, B.T. (2006a). Thyroid and pituitary gland development from hatching through metamorphosis of a teleost flatfish, the Atlantic halibut. Anat Embryol 211, 47–60.

Einarsdóttir, I.E., Silva, N., Power, D.M., Smáradóttir, H., and Björnsson, B.T. (2006b). Thyroid and pituitary gland development from hatching through metamorphosis of a teleost flatfish, the Atlantic halibut. Anat Embryol 211, 47–60.

Eldred, K.C., Hadyniak, S.E., Hussey, K.A., Brenerman, B., Zhang, P.-W., Chamling, X., Sluch, V.M., Welsbie, D.S., Hattar, S., Taylor, J., et al. (2018). Thyroid hormone signaling specifies cone subtypes in human retinal organoids. Science 362, eaau6348.

Ferraresso, S., Bonaldo, A., Parma, L., Cinotti, S., Massi, P., Bargelloni, L., and Gatta, P.P. (2013). Exploring the larval transcriptome of the common sole (Solea solea L.). BMC Genomics 14, 315.

Ferreira, J.A., and Zwinderman, A.H. (2006). On the Benjamini–Hochberg method. Ann. Statist. 34, 1827–1849.

Fraher, D., Sanigorski, A., Mellett, N.A., Meikle, P.J., Sinclair, A.J., and Gibert, Y. (2016). Zebrafish Embryonic Lipidomic Analysis Reveals that the Yolk Cell Is Metabolically Active in Processing Lipid. Cell Reports 14, 1317–1329.

Garoche, C., Boulahtouf, A., Grimaldi, M., Chiavarina, B., Toporova, L., den Broeder, M.J., Legler, J., Bourguet, W., and Balaguer, P. (2021). Interspecies Differences in Activation of Peroxisome Proliferator-Activated Receptor γ by Pharmaceutical and Environmental Chemicals. Environ. Sci. Technol. 55, 16489–16501.

Ghaddab-Zroud, R., Seugnet, I., Steffensen, K.R., Demeneix, B.A., and Clerget-Froidevaux, M.-S. (2014). Liver X receptor regulation of thyrotropin-releasing hormone transcription in mouse hypothalamus is dependent on thyroid status. PLoS One 9, e106983.

Grimaldi, A., Buisine, N., Miller, T., Shi, Y.-B., and Sachs, L.M. (2013). Mechanisms of thyroid hormone receptor action during development: Lessons from amphibian studies. Biochimica et Biophysica Acta (BBA) - General Subjects 1830, 3882–3892.

Härer, A., Torres-Dowdall, J., and Meyer, A. (2017). Rapid adaptation to a novel light environment: The importance of ontogeny and phenotypic plasticity in shaping the visual system of Nicaraguan Midas cichlid fish ( *Amphilophus citrinellus* spp.). Mol Ecol 26, 5582–5593.

Holzer, G., and Laudet, V. (2013). Thyroid Hormones and Postembryonic Development in Amniotes. In Current Topics in Developmental Biology, (Elsevier), pp. 397–425.

Holzer, G., Besson, M., Lambert, A., François, L., Barth, P., Gillet, B., Hughes, S., Piganeau, G., Leulier, F., and Viriot, L. (2017). Fish larval recruitment to reefs is a thyroid hormone-mediated metamorphosis sensitive to the pesticide chlorpyrifos. ELife 6.

Hulbert, A.J. (2021). The under-appreciated fats of life: the two types of polyunsaturated fats. Journal of Experimental Biology 224, jeb232538.

Hulbert, A.J., and Else, P.L. (2000). Mechanisms Underlying the Cost of Living in Animals. Annual Review of Physiology 62, 207–235.

Job, S., and Shand, J. (2001). Spectral sensitivity of larval and juvenile coral reef fishes: implications for feeding in a variable light environment. Mar. Ecol. Prog. Ser. 214, 267–277.

Joshi, N., and Fass, J. (2011). Sickle: A sliding-window, adaptive, quality-based trimming tool for FastQ files (Version 1.33) [Software]. Available at https://Github.Com/Najoshi/Sickle.

Kawakami, Y., Yokoi, K., Kumai, H., and Ohta, H. (2008). The role of thyroid hormones during the development of eye pigmentation in the Pacific bluefin tuna (Thunnus orientalis). Comparative Biochemistry and Physiology Part B: Biochemistry and Molecular Biology 150, 112–116.

Kim, D., Langmead, B., and Salzberg, S.L. (2015). HISAT: a fast spliced aligner with low memory requirements. Nature Methods 12, 357–360.

Kulkarni, M.M. (2011). Digital Multiplexed Gene Expression Analysis Using the NanoString nCounter System. Current Protocols in Molecular Biology 94, 25B.10.1-25B.10.17.

Laudet, V. (2011). The Origins and Evolution of Vertebrate Metamorphosis. Current Biology 21, R726– R737.

Lehmann, R., Lightfoot, D.J., Schunter, C., Michell, C.T., Ohyanagi, H., Mineta, K., Foret, S., Berumen, M.L., Miller, D.J., Aranda, M., et al. (2019). Finding nemo’s genes: A chromosome-scale reference assembly of the genome of the orange clownfish Amphiprion percula. Molecular Ecology Resources 19, 570–585.

Lowe, W.H., Martin, T.E., Skelly, D.K., and Woods, H.A. (2021). Metamorphosis in an Era of Increasing Climate Variability. Trends in Ecology & Evolution 36, 360–375.

Martin, M. (2011). Cutadapt removes adapter sequences from high-throughput sequencing reads. EMBnet.Journal 17, 10.

Mazurais, D., Darias, M., Zambonino-Infante, J.L., and Cahu, C.L. (2011). Transcriptomics for understanding marine fish larval development. Canadian Journal of Zoology 89, 599–611.

McMenamin, S.K., and Parichy, D.M. (2013). Metamorphosis in Teleosts. In Current Topics in Developmental Biology, (Elsevier), pp. 127–165.

Miao, Y., Wu, W., Dai, Y., Maneix, L., Huang, B., Warner, M., and Gustafsson, J.-Å. (2015). Liver X receptor β controls thyroid hormone feedback in the brain and regulates browning of subcutaneous white adipose tissue. Proc Natl Acad Sci U S A 112, 14006–14011.

Mitchell, L.J., Cheney, K.L., Lührmann, M., Marshall, J., Michie, K., and Cortesi, F. (2021). Molecular Evolution of Ultraviolet Visual Opsins and Spectral Tuning of Photoreceptors in Anemonefishes (Amphiprioninae). Genome Biology and Evolution 13, evab184.

Mullur, R., Liu, Y.-Y., and Brent, G.A. (2014). Thyroid hormone regulation of metabolism. Physiol Rev 94, 355–382.

Nishimura, T. (2020). Feedforward Regulation of Glucose Metabolism by Steroid Hormones Drives a Developmental Transition in Drosophila. Current Biology 30, 3624–3632.e5.

Pinto, C.L., Kalasekar, S.M., McCollum, C.W., Riu, A., Jonsson, P., Lopez, J., Swindell, E.C., Bouhlatouf, A., Balaguer, P., Bondesson, M., et al. (2016). Lxr regulates lipid metabolic and visual perception pathways during zebrafish development. Molecular and Cellular Endocrinology 419, 29–43.

Power, D.M., Einarsdóttir, I.E., Pittman, K., Sweeney, G.E., Hildahl, J., Campinho, M.A., Silva, N., Sæle, Ø., Galay-Burgos, M., Smáradóttir, H., et al. (2008). The Molecular and Endocrine Basis of Flatfish Metamorphosis. Reviews in Fisheries Science 16, 95–111.

Rennison, D.J., Owens, G.L., Heckman, N., Schluter, D., and Veen, T. (2016). Rapid adaptive evolution of colour vision in the threespine stickleback radiation. Proc. R. Soc. B. 283, 20160242.

Roux, N., Salis, P., Lambert, A., Logeux, V., Soulat, O., Romans, P., Frédérich, B., Lecchini, D., and Laudet, V. (2019). Staging and normal table of postembryonic development of the clownfish (Amphiprion ocellaris). Developmental Dynamics 248, 545–568.

Roux, N., Salis, P., Lee, S.-H., Besseau, L., and Laudet, V. (2020). Anemonefish, a model for Eco-Evo- Devo. EvoDevo 11, 20.

Roux, N., Logeux, V., Trouillard, N., Pillot, R., Magré, K., Salis, P., Lecchini, D., Besseau, L., Laudet, V., and Romans, P. (2021). A star is born again: Methods for larval rearing of an emerging model organism, the False clownfish *Amphiprion ocellaris*. J Exp Zool (Mol Dev Evol) jez.b.23028.

Russo, S.C., Salas-Lucia, F., and Bianco, A.C. (2021). Deiodinases and the Metabolic Code for Thyroid Hormone Action. Endocrinology 162, bqab059.

Sachs, L.M., and Buchholz, D.R. (2017). Frogs model man: *In vivo* thyroid hormone signaling during development: SACHS and BUCHHOLZ. Genesis 55, e23000.

Salis, P., Roux, N., Huang, D., Marcionetti, A., Mouginot, P., Reynaud, M., Salles, O., Salamin, N., Pujol, B., Parichy, D.M., et al. (2021). Thyroid hormones regulate the formation and environmental plasticity of white bars in clownfishes. Proc Natl Acad Sci USA 118, e2101634118.

Sayre, N.L., and Lechleiter, J.D. (2012). Fatty acid metabolism and thyroid hormones. Current Trends Endocrinology 1, 65–76.

Seuront, L., Schmitt, F., and Lagadeuc, Y. (2001). Turbulence intermittency, small-scale phytoplankton patchiness and encounter rates in plankton: where do we go from here? Deep Sea Research Part I: Oceanographic Research Papers 48, 1199–1215.

Shand, J. (1997). Ontogenetic changes in retinal structure and visual acuity: a comparative study of coral-reef teleosts with differing post-settlement lifestyles. Environmental Biology of Fishes 49, 307– 322.

Shand, J., Davies, W.L., Thomas, N., Balmer, L., Cowing, J.A., Pointer, M., Carvalho, L.S., Trezise, A.E.O., Collin, S.P., Beazley, L.D., et al. (2008). The influence of ontogeny and light environment on the expression of visual pigment opsins in the retina of the black bream, Acanthopagrus butcheri. Journal of Experimental Biology 211, 1495–1503.

Shao, C., Bao, B., Xie, Z., Chen, X., Li, B., Jia, X., Yao, Q., Ortí, G., Li, W., and Li, X. (2017). The genome and transcriptome of Japanese flounder provide insights into flatfish asymmetry. Nature Genetics 49, 119.

Sheridan, Mark.A., and Kao, Y.-H. (1998). Regulation of Metamorphosis-Associated Changes in the Lipid Metabolism of Selected Vertebrates. Am Zool 38, 350–368.

Sinha, R.A., Singh, B.K., and Yen, P.M. (2018). Direct effects of thyroid hormones on hepatic lipid metabolism. Nat Rev Endocrinol 14, 259–269.

Stahlberg, A., Kubista, M., and Pfaffl, M. (2004). Comparison of Reverse Transcriptase in Gene Expression Analysis. Clinical Chemistry 50, 1678–1679.

Suzuki, S.C., Bleckert, A., Williams, P.R., Takechi, M., Kawamura, S., and Wong, R.O.L. (2013). Cone photoreceptor types in zebrafish are generated by symmetric terminal divisions of dedicated precursors. Proceedings of the National Academy of Sciences 110, 15109–15114.

Tagawa, M., and Hirano, T. (1989). Changes in tissue and blood concentrations of thyroid hormones in developing chum salmon. General and Comparative Endocrinology 76, 437–443.

Tata, J.R. (2006). Amphibian metamorphosis as a model for the developmental actions of thyroid hormone. Molecular and Cellular Endocrinology 246, 10–20.

Tocher, D.R. (2010). Fatty acid requirements in ontogeny of marine and freshwater fish. Aquaculture Research 41, 717–732.

Untergasser, A., Cutcutache, I., Koressaar, T., Ye, J., Faircloth, B.C., Remm, M., and Rozen, S.G. (2012). Primer3—new capabilities and interfaces. Nucleic Acids Res 40, e115.

Wang, B., and Tontonoz, P. (2018). Liver X receptors in lipid signalling and membrane homeostasis. Nat Rev Endocrinol 14, 452–463.

Wang, Z., Guan, D., Wang, S., Chai, L.Y.A., Xu, S., and Lam, K.-P. (2020). Glycolysis and Oxidative Phosphorylation Play Critical Roles in Natural Killer Cell Receptor-Mediated Natural Killer Cell Functions. Frontiers in Immunology 11.

Xie, D., Chen, C., Dong, Y., You, C., Wang, S., Monroig, Ó., Tocher, D.R., and Li, Y. (2021). Regulation of long-chain polyunsaturated fatty acid biosynthesis in teleost fish. Progress in Lipid Research 82, 101095.

Zhang, Q., You, C., Liu, F., Zhu, W., Wang, S., Xie, D., Monroig, Ó., Tocher, D.R., and Li, Y. (2016). Cloning and Characterization of Lxr and Srebp1, and Their Potential Roles in Regulation of LC-PUFA Biosynthesis in Rabbitfish *Siganus canaliculatus*. Lipids 51, 1051–1063.

